# Neuronal histone methyltransferase EZH2 regulates neuronal morphogenesis, synaptic plasticity, and cognitive behavior of mice

**DOI:** 10.1101/582908

**Authors:** Mei Zhang, Yong Zhang, Qian Xu, Joshua Crawford, Cheng Qian, Guo-Hua Wang, Eastman Lewis, Philip Hall, Gül Dolen, Richard L. Huganir, Jiang Qian, Xin-Zhong Dong, Mikhail V. Pletnikov, Chang-Mei Liu, Feng-Quan Zhou

## Abstract

Recent studies showed that in the nervous system histone methyltransferase EZH2-mediated trimethylation of histone H3 lysine 27 (H3K27me3) acts to regulate neural stem cell proliferation and fate specificity through silencing different gene sets. Here we explored the function of EZH2 in early post-mitotic excitatory neurons by generating a neuronal specific *Ezh2* conditional knockout mouse line. The results showed that lack of neuronal EZH2 led to delayed neuronal migration, more complex dendritic arborization, and significantly increased dendritic spine density. RNA-sequencing (RNA-seq) experiments comparing control and *Ezh2* knockout neurons revealed that neuronal EZH2 regulated genes related to neuronal morphogenesis. In particular, *Pak3* was identified as a target gene suppressed by EZH2 and H3K27me3, and expression of dominant negative PAK3 reversed *Ezh2* knockout-induced higher dendritic spine density. Lastly, lack of neuronal EZH2 resulted in impaired memory behaviors in adult mice. Our results demonstrated that neuronal EZH2 played important roles in controlling multiple steps of neuronal morphogenesis during development, which had long-lasting effects on cognitive function in adult mice.

## Introduction

During the development of the central nervous system (CNS), post-mitotic neurons generated through neurogenesis undergo several steps of neuronal morphogenesis, including neuronal migration, neuronal polarization, axon growth/guidance, dendrite development, and synaptogenesis, to form the functional neural circuits. Disruptions in these processes are believed to cause many neurodevelopmental disorders, such as autism spectrum disorder (ASD), schizophrenia, and intellectual disability. Previous studies have extensively investigated the roles of extracellular guidance cues, their receptors and downstream signaling mediators, as well as cytoskeletal proteins, in regulation of these processes. Epigenetic regulation independent of DNA sequence is emerging to be a key cellular mechanism for coordinated regulation of functionally relevant gene expression in neural development. Interestingly, perturbation of several histone methyltransferases, such as *NSD1* (H3K36) and *MLL2* (H3K4), causes Sotos Syndrome and Kabuki Syndrome, respectively, both of which show intellectual disability (Douglas et al., 2003; Ng et al., 2010). It appears that these histone-modifying proteins are emerging as causes of neurodevelopmental disorders.

The polycomb repressive complex 2 (PRC2) is one of the two polycomb group proteins (PcG) that can methylate the H3 histone lysine 27 (H3K27me3) (Margueron and Reinberg, 2011). The mouse PRC2 has 4 subunits: SUZ12, EED, EZH2, and RbAp48, among which EZH2 is the methyltransferase that trimethylates H3K27. In addition to methyltransferase, H3K27 methylation is also negatively regulated by 2 demethylases, JMJD3 and UTX which remove methyl groups specifically from H3K27 (Agger et al., 2007; De Santa et al., 2007; Hong et al., 2007; Lan et al., 2007). Previous studies have identified *EZH2* as one of the causative genes of the Weaver Syndrome, a developmental disorder characterized by macrocephaly, dysmorphic facial features, accelerated skeletal maturation, limb anomalies, development delay and intellectual disability (Gibson et al., 2012; Tatton-Brown et al., 2011). In addition, a genome-wide profiling of histone methylations has shown a significant difference in H3K27 methylation between olfactory cells (with neuronal traits) obtained from the schizophrenia patients and those from control groups (Kano et al., 2012). Moreover, both *EZH2* and *JMJD3* have been identified as susceptible genes for ASD (Iossifov et al., 2012; Li et al., 2016). These facts prompted us to study the function of EZH2 and H3K27 methylation in early neuronal development of the brain.

The main function of PRC2 and H3K27 methylation is to epigenetically maintain gene silencing (Beisel and Paro, 2011). Previous studies in stem cell maintenance and differentiation suggested that PRC2 function to maintain cell identify at different stages of cell development (Hwang et al., 2014; Margueron and Reinberg, 2011; Pereira et al., 2010; Schoeftner et al., 2006). For instance, in stem cells, H3K27me3 represses the expression of differentiation-related genes to maintain stem cell fate. When stem cells differentiate into specific cell types in response to extracellular signals, H3K27me3 instead represses stem cell genes and genes related to other cell types. Thus, H3K27me3 acts to repress different sets of genes in the same cell lineage at different developmental stages. EZH2 is highly expressed in dividing neural progenitors, and loss of EZH2 in neural progenitor cells disrupts neurogenesis in both embryonic and adult animals (Pereira et al., 2010; Zhang et al., 2014). When neural progenitors differentiate into post-mitotic neurons, the expression of EZH2 is downregulated, but still maintained. Conditionally knocking out *Ezh2* in neural progenitors has been shown to regulated directional neuronal migration in the brain stem (Di Meglio et al., 2013). Similarly, knocking down EZH2 in radial glia cells led to impaired cortical neuronal migration (Zhao et al., 2015). Very interestingly, a recent study showed that activating endogenous EZH2 in embryonic stem cells using CRISPRa approach promoted neuronal differentiation, likely by suppressing the expression of endodermal and mesodermal specific gene transcription (Liu et al., 2018). Moreover, when combined with other transcription factors, such as POU3F2 and Neurogenin 1, EZH2 expression could induce direct reprogramming of fibroblasts to neurons. In sum, these studies highlighted the important roles of EZH2 in stem cells to control neuronal differentiation. However, the roles of EZH2 in post-mitotic neurons remain elusive.

As mentioned above, EZH2 in neural progenitors and in post-mitotic neurons may regulate different sets of genes and have distinct neuronal phenotypes. More importantly, to date the specific *in vivo* roles of EZH2 in early post-mitotic neurons during development remain largely unknown. To directly address the *in vivo* roles of EZH2 specifically in excitatory post-mitotic neurons, we generated neuron-specific *Ezh2* conditional knockout mice using *Neurod6-Cre* mice, in which Cre recombinase is expressed specifically in early post-mitotic excitatory neurons of the developing brain. We found that deletion of neuronal EZH2 delayed but not persistently blocked neuronal migration. More interestingly, EZH2 null neurons displayed more complex dendritic arborization and increased dendritic spine density, indicating that EZH2 acts to suppress dendritic development. RNA-Seq experiments comparing control and EZH2 null neurons identified genes regulated by EZH2, among which *Pak3* was selected for further study. The results showed that PAK3 expression was suppressed by EZH2 through H3K27me3. Functionally, expression of a dominant negative PAK3 reversed the increased dendritic spine density in EZH2 null neurons. Lastly, we provided evidence that adult mice lacking neuronal EZH2 had reduced working and recognition memory. Together, our study revealed an important role of neuronal EZH2 in regulating multiple steps of neuronal morphogenesis, which affected cognitive function in adult mice.

## Results

### Conditional deletion of EZH2 in post-mitotic pyramidal neurons during development

To study the roles of EZH2 in post-mitotic neurons, we first performed immunostaining to examine the presence of EZH2 and H3K27me3. The results showed that EZH2 and SUZ12, two major components of the PRC2 complex, and H3K27me3 were all clearly presented in the nuclei of post-mitotic cortical neurons (Supplementary Fig. S1a). To delete EZH2 specifically in post-mitotic neurons, we generated neuronal specific conditional EZH2 knockout mice by breeding *Ezh2* floxed mice (*Ezh2*^*f/f*^) with *Neurod6-Cre* mice, which express the Cre recombinase mainly in early developing post-mitotic neurons and some in intermediate progenitors (Goebbels et al., 2006). Previous studies (Goebbels et al., 2006; Wu et al., 2005) using Cre reporter mice showed that *Neurod6-Cre*-mediated recombination occurs in almost all excitatory pyramidal neurons in the dorsal telencephalon staring at about embryonic day 11 (E11). Thus, several recent studies used this mouse line to study gene functions specifically in post-mitotic neurons during development (Hirayama et al., 2012; Kazdoba et al., 2012; Morgan-Smith et al., 2014; Schwab et al., 2000). In support, immunostaining of CRE in postnatal day 0 (P0) *Neurod6-Cre* mouse brain sections showed wide spread CRE expression in cortex and hippocampus (Supplementary Fig. S1b).

Although both *Ezh2* and *Neurod6* are located in the chromosome 6, we were able to obtain a few *Neurod6-Cre*/*Ezh2*^*f/+*^ mice, which were then bred with *Ezh2*^*f/f*^ mice to obtain the conditional neuronal EZH2 knockout mice, *Neurod6-Cre*/*Ezh2*^*f/f*^ mice (named *Ezh2*^*Δ/Δ*^ mice hereafter). The CRE-mediated recombination was first verified by RT-PCR showing a CRE-generated band at 254 bp (Supplementary Fig. S1c), the same as shown in a previous study in which *Ezh2* was knocked out in B cells using the same *Ezh2*^*f/f*^ mice (Su et al., 2003). To directly show that EZH2 was deleted in post-mitotic neurons, we examined the EZH2 level in cultured E18 hippocampal neurons from control or *Ezh2*^*Δ/Δ*^ mice by immunostaining. EZH2 was clearly localized in wildtype neurons from the control mice, whereas it was drastically decreased in *Cre*-positive neurons from *Ezh2*^*Δ/Δ*^ mice (Fig. 1a, b). We next examined the fluorescence intensity of H3K27me3 immunostaining in cultured hippocampal neurons from *Ezh2*^*Δ/Δ*^ mice. The results showed that the levels of H3K27me3 were significantly decreased in cultured *Cre*-positive cells from *Ezh2*^*Δ/Δ*^ mice (Fig. 1c, d). We also examined the H3K27me3 levels in cortical slices and observed a similar significant reduction in *Cre*-positive cortical neurons from *Ezh2*^*Δ/Δ*^ mice *in vivo* (Fig. 1e, f). The expression levels of EZH2 and H3K27me3 in *Ezh2*^*Δ/Δ*^ mice were also evaluated by western blot. The western blot of postnatal day 0 (P0) mouse cortical lysate showed that EZH2 level in *Ezh2*^*Δ/Δ*^ mice was dramatically decreased compared to the *Cre*-negative control littermate mice (*Ezh2*^*f/f*^) (Fig. 1g). The remaining EZH2 was most likely from interneurons or developing glial cells that were *Cre*-negative cells. Because some studies have shown that EZH1, the paralog of EZH2, had similar methyltransferase activity (Henriquez et al., 2013), the remaining H3K27me3 in *Ezh2*^*Δ/Δ*^ neurons could also be due to the effect of EZH1. Taken together, the results demonstrated clearly that EZH2 was successfully deleted in developing cortical and hippocampal neurons of *Ezh2*^*Δ/Δ*^ mice, and deleting EZH2 alone was sufficient to significantly reduce the H3K27me3 levels in neurons in the presence of EZH1. *Ezh2*^*Δ/Δ*^ mice thus provide us a useful model to study the specific function of EZH2 and H3K27me3 in post-mitotic neurons.

**Figure 1.**
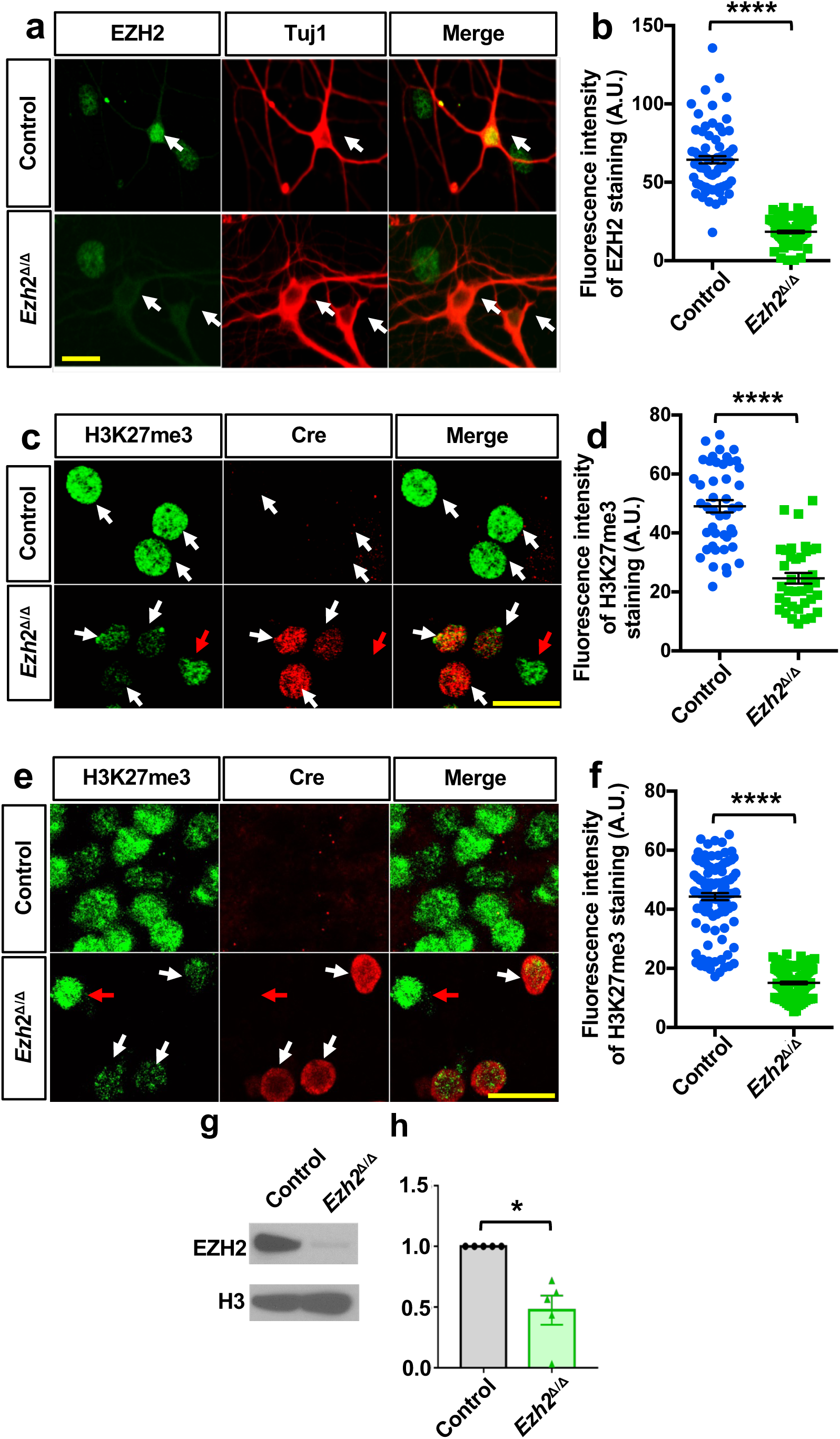
Deletion of H3K27 methyltransferase *Ezh2* in post-mitotic neurons. (a) Representative images showing the deletion of EZH2 in Cre-positive hippocampal neurons of *Ezh2*^Δ/Δ^ mice *in vitro*. detected by immunostaining of EZH2. The white arrows indicate the Tuj1 positive neurons. Scale bar, 20µm. (b) Quantification of the fluorescence intensity of EZH2 staining shown in (a). n=74 and 91 for the control and *Ezh2*^*Δ/Δ*^ neurons, respectively, from three independent experiments. \*\*\*\**P*<0.0001. (c) Representative images showing reduced level of H3K27me3 expression in Cre-positive hippocampal neurons *in vitro*. The white arrows in the upper panel indicate Cre negative neurons from the control mice. The white arrows in the bottom panel indicate the Cre positive neurons from the *Ezh2*^*Δ/Δ*^ mice, whereas the red arrow indicates a Cre negative neuron. Scale bar, 20µm. (d) Quantification of the fluorescence intensity of H3K27me3 staining shown in (c). n=45 and 37 for the control and *Ezh2*^*Δ/Δ*^ neurons, respectively, from three independent experiments. \*\*\*\**P*<0.0001. (e) Representative images showing reduced level of H3K27me3 in cortical neurons from cortical slices of *Ezh2*^Δ/Δ^ mice *in vivo* by immunostaining of H3K27me3. The white arrows in the bottom panel indicate the Cre positive neurons in the *Ezh2*^*Δ/Δ*^ cortical slices, and the red arrow indicates a Cre negative neuron. Scale bar, 20µm. (f) Quantification of the fluorescence intensity of H3K27me3 staining shown in (e). n=103 and 111 neurons for the control and *Ezh2*^*Δ/Δ*^ neurons, respectively, from three independent experiments. \*\*\*\**P*<0.0001. (g) Representative western blot images showing significant reduction of EZH2 protein levels in cortical lysate from *Ezh2*^Δ/Δ^ mice compared to control at P0. Histone 3 (H3) was used as the loading control. (h) Quantification of western blot results shown in (g) by measuring the ratio of EZH2 and H3, and the data were normalized to the control (*P* = 0.0117, n=5 mice for each condition). Data are represented as mean ± SEM. \**P*<0.05, \*\**P*<0.01, \*\*\**P*<0.001, \*\*\*\**P* < 0.0001, compared to control if not designated.

### Neuronal EZH2 regulates migration of post-mitotic cortical neurons *in vivo*

To explore neuronal functions of EZH2 during neural development, we analyzed the structure of mouse cortex in both *Ezh2*^*Δ/Δ*^ mice and their littermate *Ezh2*^*f/f*^ control mice at different developmental stages. The structure of forebrain was analyzed by Nissl staining at P0, and no significant difference in brain structure was observed in *Ezh2*^*Δ/Δ*^ mutant mice (Supplementary Fig. S2a). Because *Neurod6*-Cre is mainly expressed in post-mitotic neurons (Goebbels et al., 2006) and a small percentage of it has been identified in intermediate progenitors of the dorsal telencephalon (Wu et al., 2005), during early neuronal morphogenesis processes, such as neuronal migration, Cre is likely efficiently expressed in later born upper cortical layer neurons at that particular time point. Indeed, when the cortical lamination was assessed at E15, a time point of generation of earlier born deep layer cortical neurons, the Tbr1 positive deep layer neurons appeared to be similar between the control and the *Ezh2*^*Δ/Δ*^ mice (Supplementary Fig. S2b). The potential reason is that EZH2 protein might not been fully deleted in these early born deep layer post-mitotic neurons. To investigate the effects of neuronal EZH2 on the development of upper layer cortical neurons, the mouse cortex was analyzed at P0 by Foxp1 and Ctip2 staining, which labeled layers II-IV and V-VI, respectively (Molyneaux et al., 2007). The distributions of labeled cells within 6 bins assigned across the apicobasal axis were quantified. We noted a milder but significant reduction of Foxp1 expressing neurons in *Ezh2*^*Δ/Δ*^ mice at the upper position (Bin 5) compared with the control mice (Fig. 2a, b), whereas the total number of Foxp1 positive neurons was not significantly changed. In addition, the distributions of Ctip2 positive neurons were also significantly different between the *Ezh2*^*Δ/Δ*^ and the control mice (Fig. 2a, c). The Ctip2 positive neurons in *Ezh2*^*Δ/Δ*^ mice were at a higher density in Bin 3 than those in wild type mice (Fig. 2c). Together, these results suggested that deletion of neuronal EZH2 by *Neurod6-Cre* significantly reduced the migration of upper cortical neurons.

**Figure 2.**
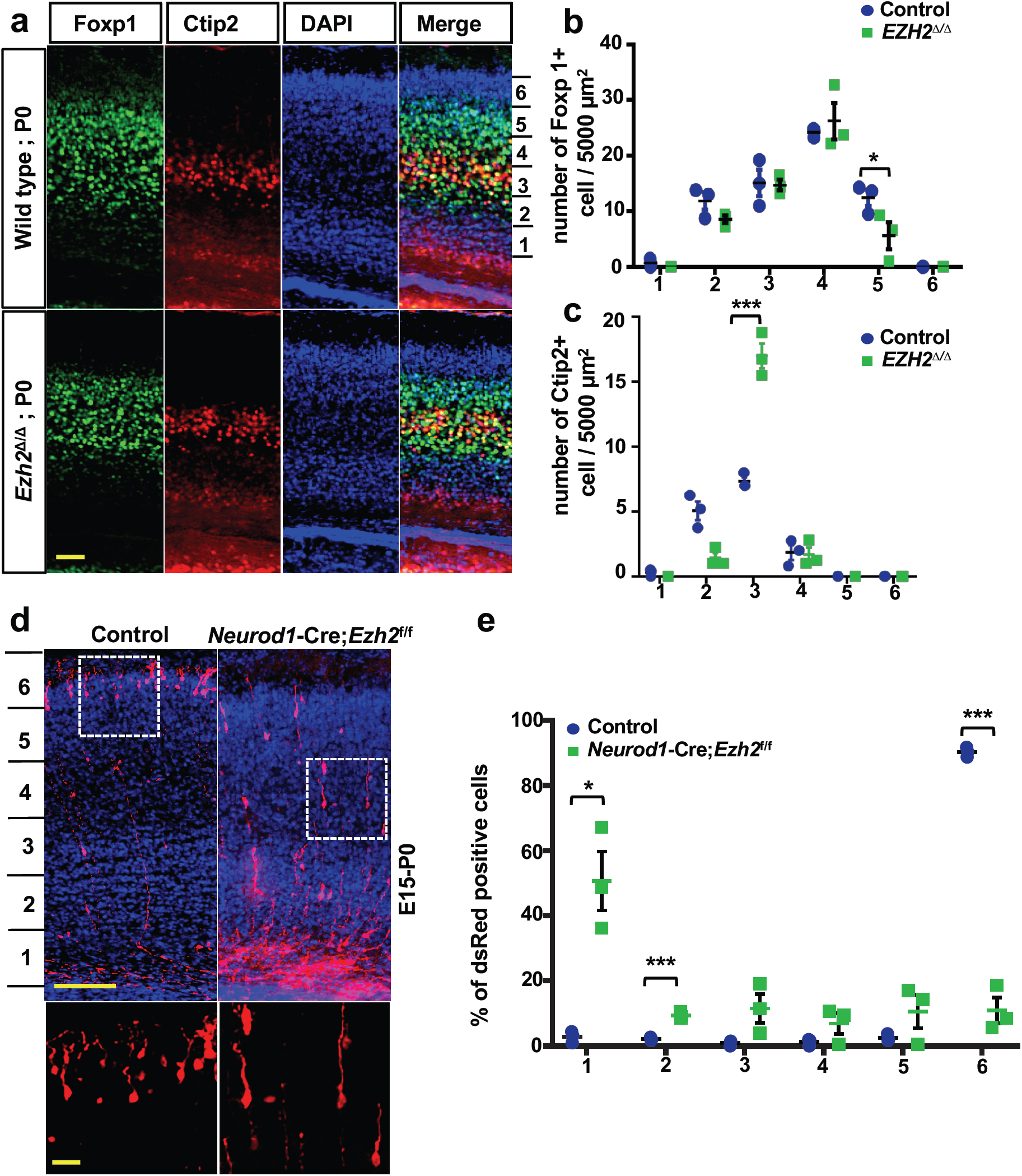
Knocking out EZH2 in cortical neurons delayed the migration of upper cortical neurons. (a) Representative images of mouse cortex immunostained for Foxp1 and Ctip2 in control and *EZH2*^Δ/Δ^ mice at P0. Foxp1 and Ctip2 labeling help to define the layer II-IV and IV-V of the cortical plate, respectively. The distributions of labeled cells were assigned into 6 bins across the apicobasal axis. Scale bar, 100 µm. (b) Quantification of Foxp1 positive cells in 6 equal bins showing significantly reduced Foxp1 positive neurons in Bin 5 (Bin1, *P*=0.2495; Bin2, *P*=0.1263; Bin3, *P*=0.9192; Bin4, *P*=0.5790; Bin5, *P*=0.0213, n=3 mice for each condition). (c) Quantification of Ctip2 positive cells in 6 equal bins showing significantly increased Ctip2 positive neurons in Bin 3 (Bin1, *P*=0.1885; Bin2, *P*=0.0822; Bin3, *P*=0.016; Bin4, *P*=0.8954, n=3 mice for each condition). (d) Representative confocal images of *EZH2*^*f/f*^ mice mouse cortices in utero electroporated with dsRed or dsRed/*Neurod1-Cre*. The electroporation was done at E15 and pups were sacrificed at P0 for analysis. The two white dashed line boxes in the upper panels were enlarged and presented in the bottom panels. Scale bar, 200 µm in the upper panels and 50 µm in the bottom panels. (e) Quantification of dsRed positive cells in mouse cortices of (d) showing significantly increased number of cells at Bin1 but reduced number of cells at Bin6 in mice electroporated with Neurod1-Cre (Bin1, *P*=0.0325; Bin2, *P*=0.0019; Bin3, *P*=0.1375; Bin4, *P*=0.2140; Bin5, *P*=0.2502; Bin6, *P*=0.0016; n=3 mice for each condition). Data are presented as mean ± SEM. \**P*<0.05, \*\**P*<0.01, \*\*\**P*<0.001, \*\*\*\**P* < 0.0001, compared to control if not designated.

To verify if EZH2 regulates the migration of post-mitotic neurons in a cell autonomous manner and to visualize the morphologies of migrating neurons, we performed acute EZH2 deletion by in utero electroporation of *Neurod1-Cre* plasmid into E15 embryonic brains of *Ezh2*^*f/f*^ mice. The pCAG-dsRed plasmid was co-electroporated with *Neurod1-Cre* to label the transfected cells. The *Ezh2*^*f/f*^ embryos in the control group were only electroporated with the pCAG-dsRed plasmid. When the distribution of dsRed positive cells was analyzed at P0, we found that acutely knocking out EZH2 in post-mitotic neurons significantly impaired neuronal migration (Fig. 2d). In contrast, the majority of neurons in the control mice migrated into the upper cortical plate (Fig. 2d). Quantification revealed that deletion of EZH2 in post-mitotic neurons led to the migration defects of upper layer neurons at P0 (Fig. 2d, e). Neurons transition from a multipolar to a unipolar/bipolar morphology to initiate radial migration (Rakic, 1972). When neuronal morphology was examined, both control and *Ezh2*^*Δ/Δ*^ neurons showed similar polarized shape with a leading process directed toward the pial surface (Fig. 2f). When embryos were in utero electroporated at E15 and analyzed at P7, both control and *Ezh2*^*Δ/Δ*^ neurons (dsRed positive) reached the upper layer similarly (Supplementary Fig. S3). Thus, specific neuronal deletion of EZH2 only delayed but not persistently impaired the migration of cortical pyramidal neurons.

### Loss of neuronal EZH2 results in increased dendritic complexity *in vitro* and *in vivo*

To investigate the role of EZH2 in regulating neuronal morphogenesis, we isolated and cultured E18 hippocampal neurons from both control and *Ezh2*^*Δ/Δ*^ mice. At DIV5, the neurons were fixed and stained with both anti-Cre and anti-MAP2 antibodies. To quantify dendritic morphology, we first evaluated dendritic branching using the Sholl analysis, which measures the number of intersections of dendrites within each sphere plotted against radius increasing in 10µm increments of cell soma. Compared with control neurons, *Ezh2*^*Δ/Δ*^ neurons showed more complex dendritic morphology (Fig. 3a, b). The Sholl analysis of dendritic arborization revealed markedly increased dendritic branching in *Ezh2*^*Δ/Δ*^ neurons compared to those of control neurons (Fig. 3c). Further, morphological analysis revealed that the *Ezh2*^*Δ/Δ*^ neurons displayed significantly increased total dendritic branch length (TDBL) (Fig. 3d) and total dendritic branch tip number (TDBTN) (Fig. 3e). The average dendritic branch length (ADBL) was significantly reduced (Fig. 3f), primarily due to the increased proportion of short branches. These results indicated that *Ezh2*^*Δ/Δ*^ hippocampal pyramidal neurons increased their dendritic growth and arborization by promoting the formation of short dendritic branches.

**Figure 3.**
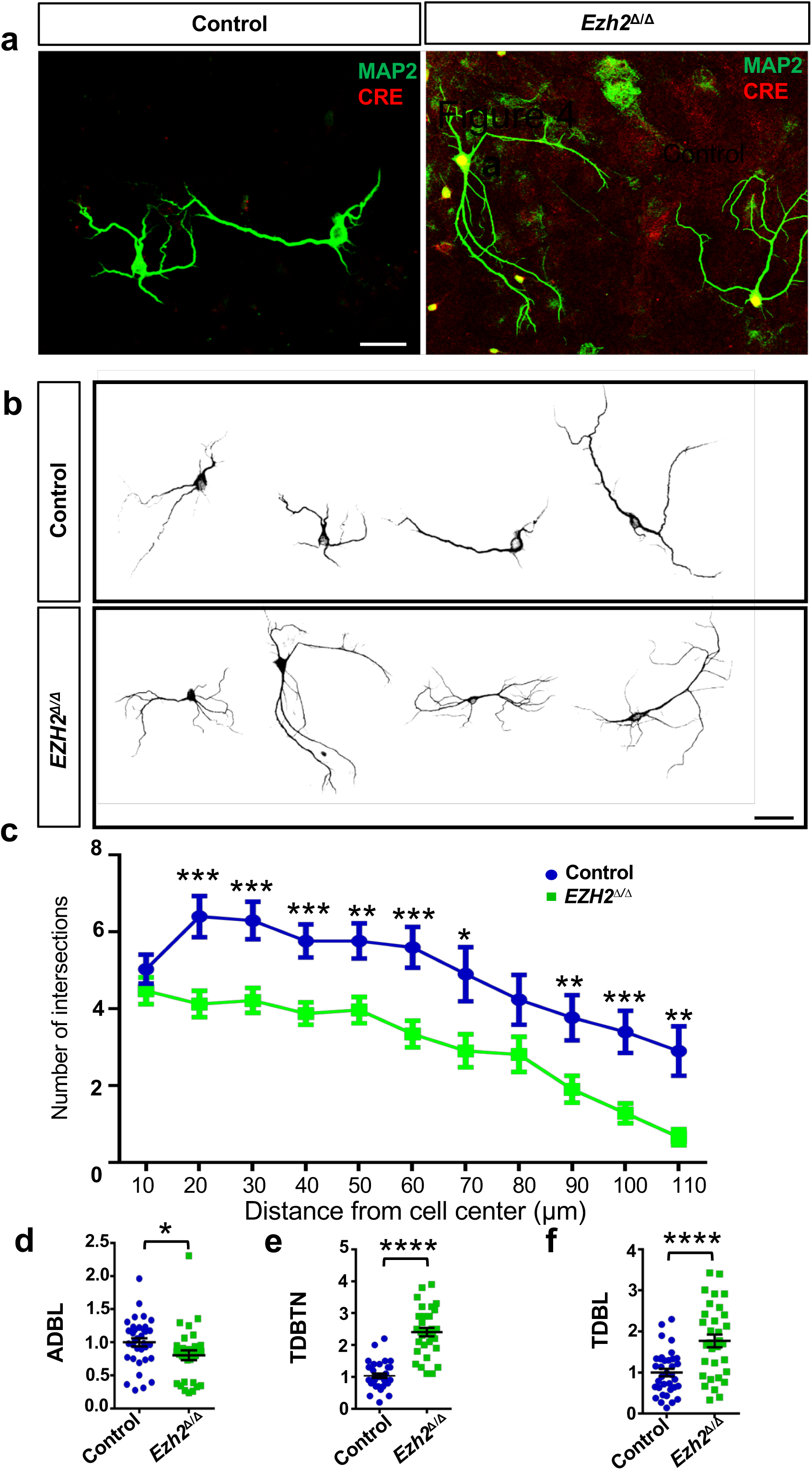
Loss of EZH2 increased dendritic branching *in vitro*. (a) Representative images of cultured hippocampal neurons from control and *Ezh2*^*Δ/Δ*^ mice stained with Cre and MAP2 antibodies. Scale bar, 20 µm. (b) Representative images of several MAP2 stained hippocampal neurons from control and *Ezh2*^*Δ/Δ*^ mice. Note the increased dendritic branching of neurons from the *Ezh2*^*Δ/Δ*^ mice. Scale bar, 20 µm. (c) Sholl analysis of hippocampal neurons from control and EZH2^Δ/Δ^ mice showing that, compared with that of the control mice, the neurons of the EZH2^Δ/Δ^ mice had increased dendritic branching. (*P*=0.2759 at 10µm; *P*=0. 0.0006 at 20 µm; *P*=0.0006 at 30 µm; *P*=0.0004 at 40 µm; *P* = 0. 0.0023 at 50 µm; *P* = 0.0006 at 60 µm; *P* = 0.0174 at 70 µm; *P* = 0.0757 at 80 µm; *P* = 0.0078 at 90 µm; *P* = 0.0007 at 100 µm; *P* = 0.0012 at 110 µm; n=30 and 32 neurons of control and *Ezh2*^*Δ/Δ*^ mice, respectively, each from 3 to 4 mice). (d) Quantification of normalized average dendritic branch length (ADBL) of control and EZH2^Δ/Δ^ neurons (*P* = 0.0441, n = 35 and 32 neurons from the control and the *Ezh2*^*Δ/Δ*^ mice, respectively, from 3 independent experiments). (e) Quantification of normalized total dendritic branch tip number (TDBTN) of control and EZH2^Δ/Δ^ neurons (*P* < 0.0001, n = 35 and 32 neurons from the control and the *Ezh2*^*Δ/Δ*^ mice, respectively, from 3 independent experiments). (f) Quantification of normalized total dendritic branch length (TDBL) of control and EZH2^Δ/Δ^ neurons *(P*<0.0001, n = 35 and 32 neurons from the control and the *Ezh2*^*Δ/Δ*^ mice, respectively, from 3 independent experiments). Data are presented as mean ± SEM. \**P*<0.05, \*\**P*<0.01, \*\*\**P*<0.001, \*\*\*\**P* < 0.0001, compared to control if not designated.

To verify if the dendritic morphology was altered in *Ezh2*^*Δ/Δ*^ mice *in vivo*, we generated *Ezh2*^*Δ/Δ*^*;Thy1-GFP-M* mice, in which pyramidal neurons in adult mouse cortex and hippocampus were labeled by Thy1-GFP. The control mice were *Ezh2*^*f/f*^*;Thy1-GFP-M* littermate mice. Because GFP labeled neurons in the hippocampus were at a higher density, making it difficult to quantify the dendritic tree clearly. We thus prepared the coronal brain sections of these mice at P28-30 and imaged the GFP-labeled layer IV-V neurons that were more sparsely labeled (Fig. 4a). Similar to cultured hippocampal neurons, *Ezh2*^*Δ/Δ*^ neurons exhibited more complex dendritic morphologies (Fig. 4b, c). Indeed, Sholl analyses revealed that *Ezh2*^*Δ/Δ*^ cortical neurons had more complex basal, but not apical, dendritic morphologies (Fig. 4d-f). Additional quantification showed that the basal dendrites of *Ezh2*^*Δ/Δ*^ neurons had significantly increased TDBTN and reduced ADBL, indicating increased dendritic branching with shorter branch lengths (Fig. 4g, h). These results were the same as those observed in cultured hippocampal neurons. However, unlike that *in vitro* results, the TDBL between the *Ezh2*^*Δ/Δ*^ and control neurons were similar (Fig. 4i). Overall, these results provided strong evidence that neuronal EZH2 plays an important role in the regulation of dendritic development.

**Figure 4.**
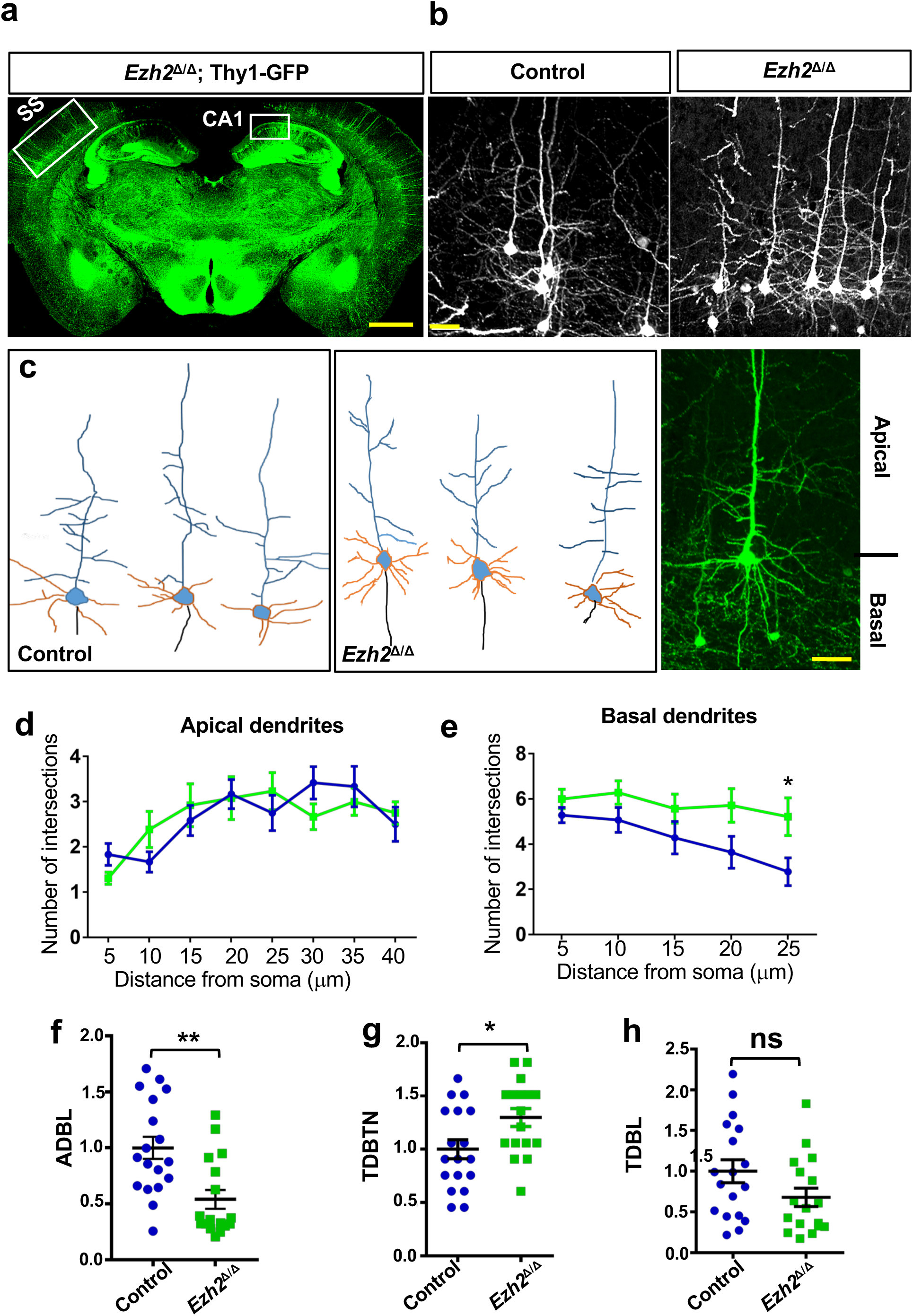
Lack of EZH2 increased the basal dendritic branching *in vivo*. (a) Representative images of *Ezh2*^*Δ/Δ*^; Thy1-GFP mice. GFP expression was detected at cortical and hippocampal region. SS indicates the somatosensory cortex and CA1 indicates the CA1 region of the hippocampus. Scale bar, 1 mm. (b) Representative images of cortical neurons in *Ezh2*^*f/f*^; Thy1-GFP and *Ezh2*^*Δ/Δ*^; Thy1-GFP mice. Scale bar, 20µm (c) Representative drawing of neurons in *Ezh2*^*f/f*^; Thy1-GFP (left panel) and *Ezh2*^*Δ/Δ*^; Thy1-GFP (middle panel) mice. The orange colored are basal dendrites and the blue colored are apical dendrites. The right panel shows a representative image of a GFP labeled cortical neuron. Scale bar, 20µm (d) Sholl analysis for apical dendrites of cortical neurons showing no significant difference in dendritic trees between control and *Ezh2*^*Δ/Δ*^ cortical neurons (*n* = 14 neurons from 4 mice for each condition). (e) Sholl analysis for basal dendrites of cortical neurons showing significantly increased complexity of *Ezh2*^*Δ/Δ*^ cortical neurons, compared with that of control mice (*P* = 0.03 at 25µm, n=14 neurons from 4 mice for each condition). (f) Quantification of normalized average dendritic branch length (ADBL) of control and *Ezh2*^*Δ/Δ*^ neurons *in vivo* (*P*=0.0014, n=18 and 17 neurons for the control and the *Ezh2*^*Δ/Δ*^ mice, respectively, from 4 mice from each condition). (g) Quantification of normalized total dendritic branch tip number (TDBTN) of control and *Ezh2*^*Δ/Δ*^ neurons *in vivo* (*P*=0.0211, n=18 and 17 neurons for the control and the *Ezh2*^*Δ/Δ*^ mice, respectively, from 4 mice from each condition). (h) Quantification of normalized total dendritic branch length (TDBL) of control and *Ezh2*^*Δ/Δ*^ neurons *in vivo* (*P*=0.0879, n=18 and 17 neurons for the control and the *Ezh2*^*Δ/Δ*^ mice, respectively, from 4 mice from each condition). Data are presented as mean ± SEM. \**P*<0.05, \*\**P*<0.01, \*\*\**P*<0.001, \*\*\*\**P* < 0.0001, compared to control if not designated.

### Neuronal EZH2 regulates dendritic spine density and synaptic function

Synaptic remodeling in the hippocampus is a key process for controlling working memory and novelty recognition (Bourne and Harris, 2007; Compte et al., 2000). Therefore, we examined dendritic spine development in hippocampal and cortical pyramidal neurons of young adult control or *Ezh2*^*Δ/Δ*^ mice. We first examined P28-30 pyramidal neurons in the hippocampal CA1 region, the spine density of the basal dendrites was significantly increased (Fig. 5a and 5b). Dendritic spines often show a myriad of morphologies, which usually correlate with different synaptic properties. For instance, large mushroom like spines usually form strong and persistent connections, whereas thin spines are more dynamic and have weaker connections. To better understand how EZH2 regulates dendritic spine development, we categorized the spine morphologies into 4 different groups, stubby spine, filopodia like spines, mushroom spine, and long thin spine (Fig. 5c). Analysis of spine morphologies on apical dendrites of CA1 neurons revealed that *Ezh2*^*Δ/Δ*^ neurons had decreased percentage of mushroom spines and increased percentage of filopodia like spines (Fig. 5d). To investigate the functional consequences of increased dendritic spine density in *Ezh2*^*Δ/Δ*^ mice, we performed whole cell patch-clamp recordings of cultured hippocampal neurons *in vitro*. mEPSC analysis revealed a reduction in mEPSC frequency in *Ezh2*^*Δ/Δ*^ neurons (Fig. 5e). However, no significant change was detected in the mEPSC amplitude. Change in mEPSC amplitude are consistent with the density/conductance of postsynaptic receptors at individual synapses. Change in mEPSC frequency is usually interpreted as change in either the presynaptic release probability at existing sites or in the number of functional synaptic sites. Here, since *Ezh2*^*Δ/Δ*^ neurons exhibited more synapses, the reduction in mEPSC frequency in *Ezh2*^*Δ/Δ*^ mutant neurons is most likely due to the silencing of excitatory synapses.

**Figure 5.**
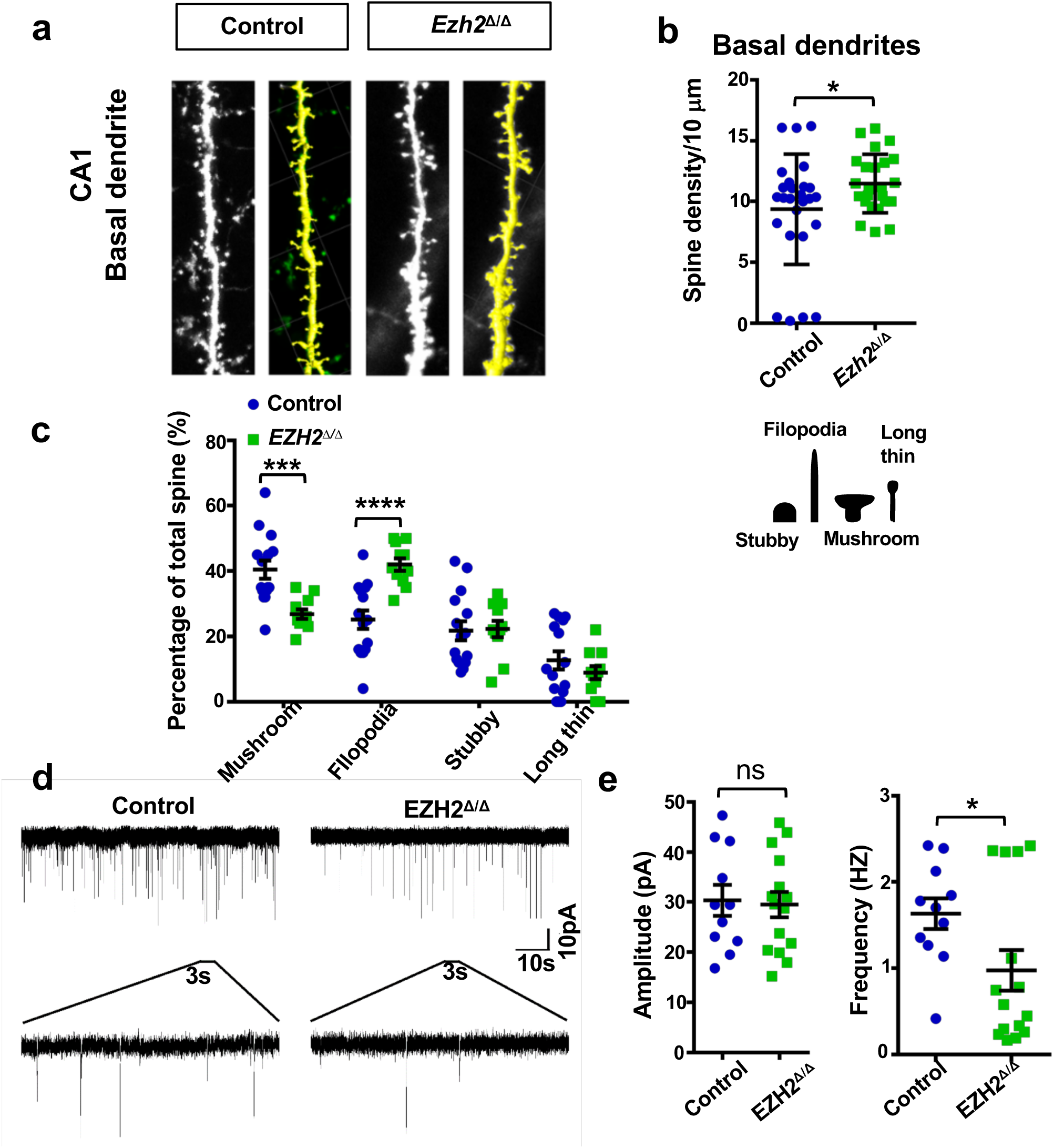
Spine density was increased in CA1 region of *EZH2*^Δ/Δ^ animals. (a) Representative images of increased spine density in basal region of hippocampal neurons from control and *Ezh2*^*Δ/Δ*^ mice. The yellow images are white images processed with the software iMaris. (b) Quantification of spine density in the basal region of hippocampal neurons from control and *Ezh2*^*Δ/Δ*^ mice (*P*=0.0260, n=28 and 24 neurons for the control and the *Ezh2*^*Δ/Δ*^ mice, respectively, from 4 mice for each condition). (c) Left: quantification analysis of different spine types in basal region of hippocampal neurons from control and *Ezh2*^*Δ/Δ*^ mice (*P*=0.0003 for mushroom; *P*=6.86624E-05 for filophodia; n=15 and 11 neurons for the control and the *Ezh2*^*Δ/Δ*^ mice, respectively, from 4 mice for each condition. Right: diagram showing 4 different spine types. (d) Representative traces of mEPSCs of hippocampal neurons from control and *Ezh2*^*Δ/Δ*^ mice (e) Quantification of average frequency and amplitude of mEPSCs of hippocampal neurons from control and *Ezh2*^*Δ/Δ*^ mice (Frequency: *P*=0.0348; Amplitude: *P*=0.8299, n=11 and 15 neurons for the control and the *Ezh2*^*Δ/Δ*^ mice, respectively, from 4 independent experiments for each condition). Data are presented as mean ± SEM. \**P*<0.05, \*\**P*<0.01, \*\*\**P*<0.001, \*\*\*\**P* < 0.0001, compared to control if not designated.

We next examined the dendritic spine density in layer IV-V cortical neurons from P28-30 *Ezh2*^*Δ/Δ*^*;Thy1*-GFP-M mice or *Ezh2*^*f/f*^*;Thy1*-GFP-M littermate mice. The spine density of basal dendrites was similarly increased in *Ezh2*^*Δ/Δ*^ mice (Supplementary Fig. S4a, b). We also performed Golgi staining in brain sections of P28-30 control or *Ezh2*^*Δ/Δ*^ mice. We found that the dendritic spine density of *Ezh2*^*Δ/Δ*^ cortical neurons was significantly increased as well compared with that of the control neurons (Supplementary Fig. S4c, d). These results indicated that EZH2 restricted dendritic spine formation, and deleting EZH2 leads to increased dendritic spine density with less mature spines.

### Transcriptome analysis of genes regulated by neuronal EZH2

To gain further insight into how EZH2 regulates gene expression in post-mitotic neurons, we examined the transcriptome between the control and *Ezh2*^*Δ/Δ*^ neurons. In addition to *Ezh2*^*Δ/Δ*^ mice, we also generated *Nestin-Cre/Ezh2*^*f/f*^ mice, in which *Ezh2* was knocked out in neural progenitors. Hippocampal neurons from E18 *Ezh2*^*f/f*^, *Ezh2*^*Δ/Δ*^ or *Nestin-Cre*/*Ezh2*^*f/f*^ embryos were dissociated and cultured in a neuronal culture condition for 3 days. The neurons were collected to extract total RNAs, which were used for RNA-seq to identify differently expressed genes (DEGs) between control and *Ezh2* knockout neurons (Fig. 6a). Our analysis revealed 1880 and 1783 up and down regulated genes, respectively, in the *Ezh2*^*Δ/Δ*^ neuronal samples compared with the control sample (fold>2). For the sample from the *Nestin-Cre*/*Ezh2*^*f/f*^ neurons, there were 2061 and 3044 up and down regulated genes, respectively (Fig. 6b, Data set 1). The DEGs identified through *Ezh2*^*Δ/Δ*^ mice and *Nestin-Cre/Ezh2*^*f/f*^ mice partially overlap. Although the RNA-seq experiments were both performed using post-mitotic neurons, this result suggested that knocking out *Ezh2* in neural progenitors or post-mitotic neurons resulted in different changes of gene transcription in neurons. To better understand the potential functions of these DEGs regulated by EZH2, we performed gene ontology (GO) analysis of biological processes. The results showed that many DEGs induced by deleting neuronal EZH2 were related to biological processes involved in neural development, such as neuron migration, positive regulation of cell migration, neuron projection morphogenesis, regulation of neuron differentiation, synapse organization, and dendrite development, etc. (Fig. 6b). We also performed network analyses of subgroups of enriched GO terms. The relationships among different GO terms were plotted and visualized using the Metascape (http://metascape.org) and Cytoscape5 (Supplementary Fig. S5a-d). Moreover, by comparing DEGs induced by EZH2 knockout with the SFARI AutDB gene database (https://gene.sfari.org), we found that many DEGs overlapped with ASD risk genes (Fig. 6c). This result is consistent with recent studies, in which EZH2 was identified to be involved in genetic etiology of autism (Li et al., 2016) or an important downstream target of CHD8, a major ASD-associated gene (Durak et al., 2016).

**Figure 6.**
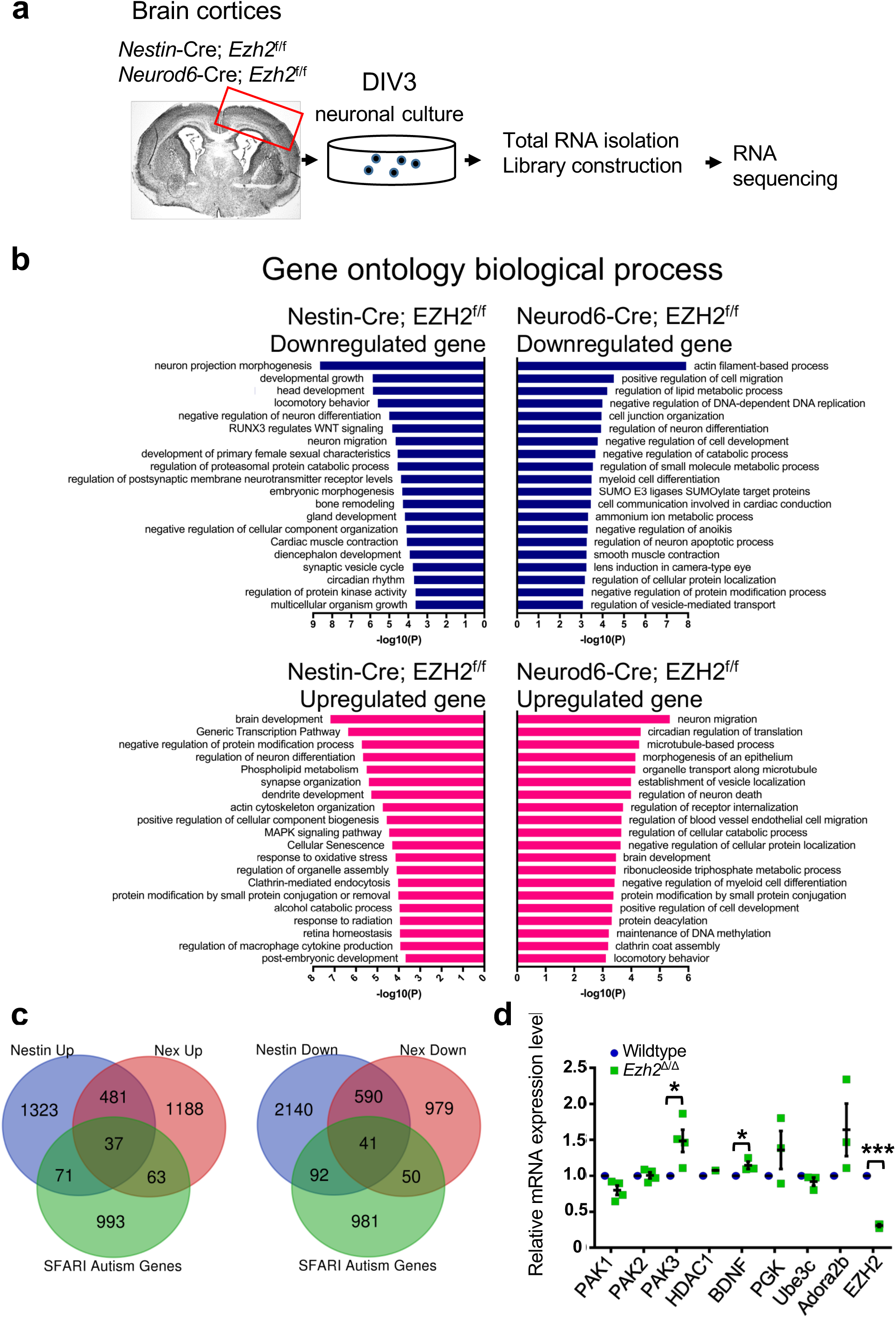
RNA-seq analysis of gene transcription in *Nestin-Cre; Ezh2*^*f/f*^ and *EZH2*^Δ/Δ^ neurons. (a) Workflow of collecting excitatory neurons from control, *Nestin-Cre; Ezh2*^*f/f*^, and *EZH2*^Δ/Δ^ mice for RNA-seq analysis. (b) Gene ontology analysis of upregulated and downregulated differential expressed genes in *Nestin-Cre; Ezh2*^*f/f*^ and *EZH2*^Δ/**Δ**^ mice, compared to that of the control mice. (c) Venn diagram representing the number of DEGs between SFARI autism genes and *Nestin-Cre; Ezh2*^*f/f*^ or *EZH2*^Δ/Δ^ mice. (d) Quantitative Reverse Transcription PCR analysis of selected DEGs in excitatory neurons in wildtype and *EZH2*^Δ/Δ^ mice (*P* = 0.02 for PAK3, *P* = 0.02 for BDNF, *P* = 0.0007 for EZH2, respectively, n=3 independent experiments). Data are presented as mean ± SEM. \**P*<0.05, \*\**P*<0.01, \*\*\**P*<0.001, \*\*\*\**P* < 0.0001, compared to control if not designated.

Lastly, using quantitative real time PCR (qPCR), we next examined a subset of genes that have been shown to be involved in regulation of dendritic development or synaptic function. Several of these genes showed significant up regulation in *Ezh2*^*Δ/Δ*^ neurons, including *Pak3* and *Bdnf* (Fig. 6d). Together, these data suggest that neuronal EZH2 plays important roles in the regulation of gene transcriptions related to neural development.

### PAK3 acts downstream of EZH2 and H3K27me3 to regulate dendritic spine development

To provide evidence that neuronal EZH2 regulates dendritic development through its downstream targeted genes, we selected PAK3 that was well-known to regulate dendritic synaptogenesis (Boda et al., 2008; Node-Langlois et al., 2006). In cultured hippocampal neurons, the protein levels of EZH2 decreased in time, whereas the levels of PAK3 increased (Fig. 7a). The result was consistent with the role of EZH2 in the regulation of H3K27me3, which functioned to repress gene expression. In supporting the qPCR result, western blot analysis showed that the protein levels of PAK3 were significantly increased in *Ezh2*^*Δ/Δ*^ neurons (Fig. 7b). We performed ChIP to determine if H3K27me3 directly interacts with the promoter region of *Pak3*. The genomic region of *Pak3* was defined into 3 overlapping fragments (R1-3, Fig. 7c), and specific primers were designed to amplify these 3 regions of *Pak3* promoter. The result showed that H3K27me3 specifically interacted with the R3 region (from 2 kb to 1 kb upstream) but not the R1/2 regions in P7 mouse cortical tissues (Fig. 7d). In contrast, immunoprecipitation with IgG did not display any interactions with R1-3 regions. The qRT-PCR data showed that the binding of H3K27me3 with the R3 region was significantly higher than that with IgG in mouse cortical tissue (Fig. 7e). These results indicated that EZH2 repressed *Pak3* expression through H3K27me3, which directly binds to the regulatory sequence upstream of *Pak3*.

**Figure 7.**
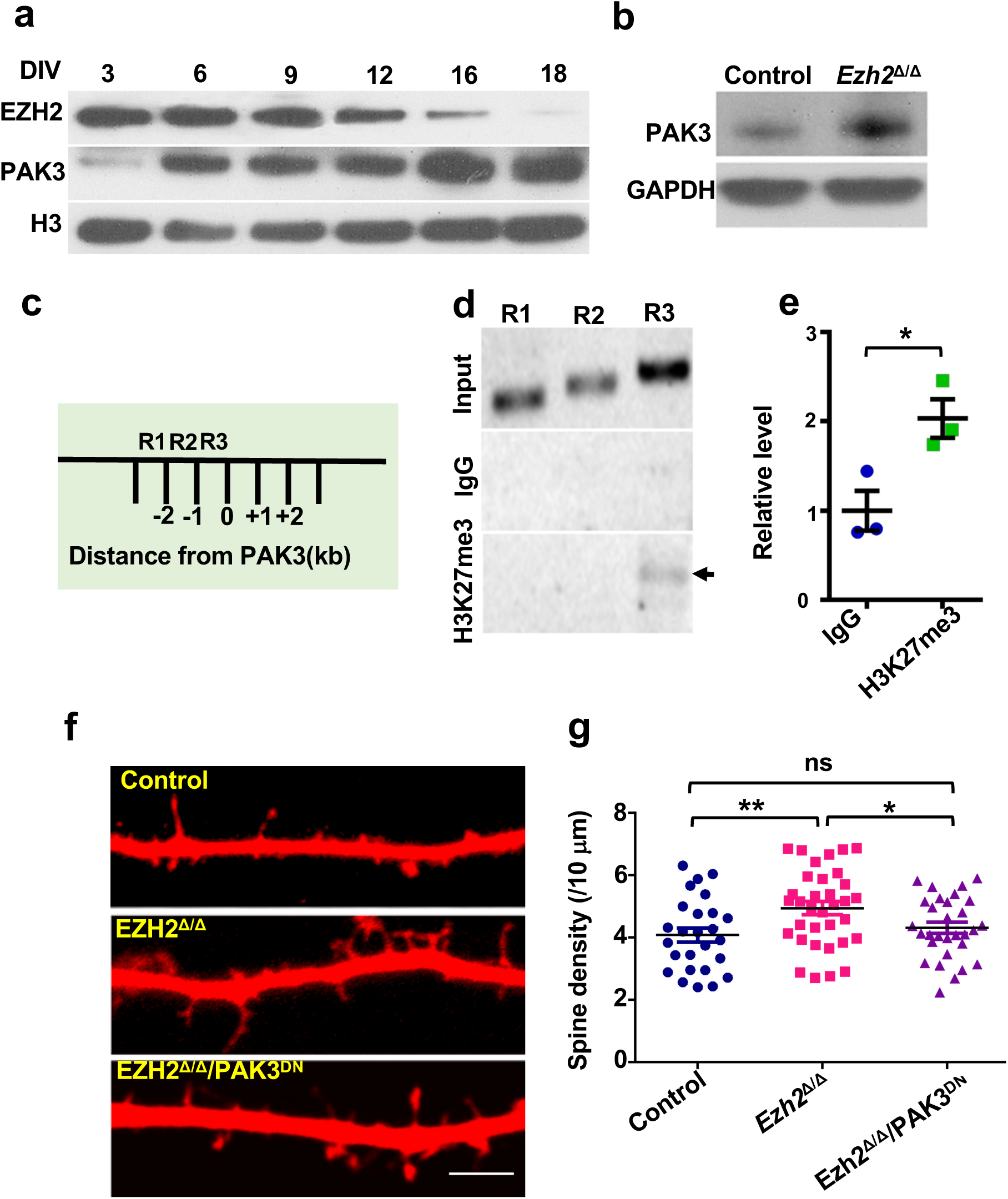
EZH2 regulates dendritic spine development by repressing gene PAK3. (a) Representative western blot images showing the expression of EZH2 and PAK3 in hippocampal neurons after different period of culturing. (b) Representative western blot images showing increased level of PAK3 in neuronal lysis from *EZH2*^Δ/Δ^ mice, compared to that from control mice. (c) Diagram showing the PAK3 promoter region, which is artificially separated into three regions, R1 to R3. (d) Representative PCR gel images showing H3K27me3 antibody associated ChIP analysis using P7 mouse cortical tissues. (e) Quantitative reverse transcription PCR analysis of the H3K27me3 ChIP assay shown in (d) (P=0.029, n=3 independent experiments). (f) Representative images of dendritic spines of cultured control neurons, *EZH2*^Δ/Δ^ neurons, and *EZH2*^Δ/Δ^ neurons expressing dominant negative PAK3 (PAK3^DN^). Note that repression of PAK3 function was able to reduced the increased spine density in *EZH2*^Δ/Δ^ neurons. Scale bar, 5µm (g) Quantification of spine densities shown in (f) (n=26, 31, and 30 neurons for the control, *EZH2*^Δ/Δ^ and PAK3^DN^ neurons, respectively, from 4 independent experiments. *P*=0.0079 between control and *EZH2*^Δ/Δ^ neurons; *P* = 0.0293 between *EZH2*^Δ/Δ^ and PAK3^DN^ neurons). Data are presented as mean ± SEM. n.s., no significant difference, \**P*<0.05, \*\**P*<0.01, \*\*\**P*<0.001, \*\*\*\**P* < 0.0001, compared to control if not designated.

To determine if PAK3 is indeed functionally involved in spine development downstream of EZH2, we tested if a dominant negative PAK3 plasmid, PAK3^K297R^ (Zhang et al., 2005), could revert the increased spine density observed in *Ezh2*^*Δ/Δ*^ neurons. E18 hippocampal neurons from the control or *Ezh2*^*Δ/Δ*^ mice were cultured. At DIV 7, the control neurons were transfected with dsRed and the *Ezh2*^*Δ/Δ*^ neurons were transfected with either dsRed or dsRed+PAK3^K297R^. At DIV18, the neurons were fixed and the dendritic spine densities were quantified. The results showed that the increased spine density in *EZH2*^*Δ/Δ*^ neurons was reverted by *Pak3* ^K297R^ to the same level as that of the control neurons (Fig. 7f, g). Altogether, these results provided strong evidence that EZH2 repressed the expression of PAK3, which regulated the spine density in hippocampal neurons.

### Neuronal EZH2 ablation during development has prolonged effects on impairing cognitive behaviors in adult mice

To investigate the potential effects of neuronal EZH2 ablation on cognitive function of adult mice, we performed a battery of behavioral tests using *Ezh2*^*Δ/Δ*^ and control (*Ezh2*^*f/f*^) littermate male and female mice. Spatial working memory and spatial recognition memory was evaluated using Y maze assay as previously described (Fig. 8a) (Ayhan, 2011; Abazyan, 2010; Dellu et al., 2000). To test for working memory, the spontaneous alternations test was performed. We found that *Ezh2*^*Δ/Δ*^ mutant mice showed a significant reduction in the percentage of correct alterations compared to control mice (Fig. 8b). Mice were also tested in the Y-maze for recognition memory (Fig. 8c). We found that *Ezh2*^*Δ/Δ*^ mice showed shorter spending time in the novel arm (the arm that was first blocked) (Fig. 8c). These results indicate that *Ezh2*^*Δ/Δ*^ mice had reduced spatial working and recognition memories. We next performed the novel object recognition test (Fig. 8a, d). The results showed that *Ezh2*^*Δ/Δ*^ mutant mice spent significantly less time sniffing the novel object (Fig. 8d), indicating that novel object recognition memory was also impaired in the *EZH2*^*Δ/Δ*^ mutant mice.

**Figure 8.**
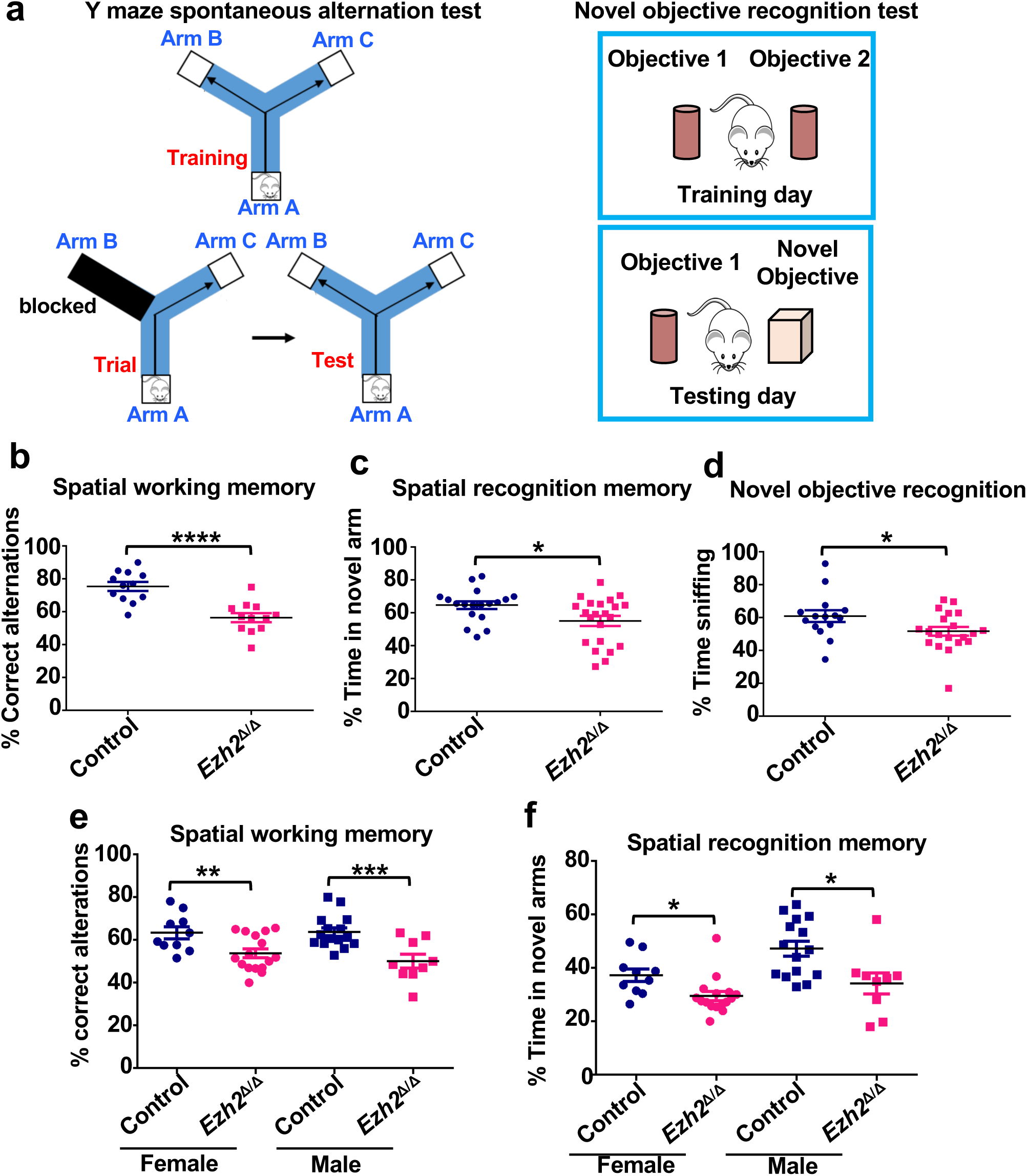
*EZH2*^*Δ/Δ*^ mutant mice deficient working and recognition memory. (a) Left: diagram of Y maze spontaneous alternation test for examining spatial working memory and spatial recognition memory. Right: diagram of the novel objective recognition test. (b) Quantification of the percentage of correct alternation between Y maze arms. The result revealed that *EZH2*^*Δ/Δ*^ mutant mice have impaired spatial working memory. *P*=3.02523E-05, n=12 for either EZH2^Δ/Δ^ or control mice. (c) Quantification of the percentage of time spent in the novel arm relative to the total duration of visits in the three arms during the test phase. Note that *EZH2*^*Δ/Δ*^ mutant mice showed reduced recognition memory. n=25 for control mice and n=18 for *EZH2*^*Δ/Δ*^ mice, *P*=0.0174. (d) Quantification of novel object recognition memory in *EZH2*^*Δ/Δ*^ mutant mice. Note that *EZH2*^*Δ/Δ*^ mice spent shorter percentage time in novel object. n=15 for control mice and n=20 for *EZH2*^*Δ/Δ*^ mice, *P*=0.0423. (e) Quantification of the percentage of alternation between Y maze arms. The result revealed that both male and female *EZH2*^*Δ/Δ*^ mutant mice have impaired short-term working memory. Female: n=10 for control mice and n=16 for *EZH2*^*Δ/Δ*^ mice, *P*=0.01; Male: n=15 for control mice and n=9 for *EZH2*^*Δ/Δ*^ mice, *P* =0.001. (f) Quantification of the percentage of time spent in novel arms. The result revealed that both male and female *EZH2*^*Δ/Δ*^ mice have impaired short-term working memory. Female: n=10 for control mice and n=16 for *EZH2*^*Δ/Δ*^ mice, *P* =0.0265; Male: n=15 for control mice and n=9 for *EZH2*^*Δ/Δ*^ mice, *P* =0.0251. Data are presented as mean ± SEM. n.s., no significant difference, \**P*<0.05, \*\**P*<0.01, \*\*\**P*<0.001, \*\*\*\**P* < 0.0001, compared to control if not designated.

Using a different group of mice, we performed similar spatial working and recognition memory tests by separately analyzing the results from male or female mice. We found that both male and female *Ezh2*^*Δ/Δ*^ mice had significantly impaired spatial working and recognition memories (Fig. 8e, f). These cognitive deficits could not be explained by changes in anxiety or general activity as we observed no group-differences in elevated plus maze test (Supplementary Fig. S6a, b) or in open field test (Supplementary Fig. S6c-e).

Collectively, these results illustrated that working and recognition memory was significantly impaired in the *Ezh2*^*Δ/Δ*^ mice, suggesting that neuronal EZH2 might be involved in neuronal development processes contributing to memory related cognitive functions in adult mice.

## Discussion

The major function of EZH2 is to trimethylate H3K27 and repress gene expression, which has been mostly studied in dividing cells, such as stem cells and cancer cells. During different stages of cell development, including the maintenance of the stem cell states and distinct cell fate differentiation, EZH2-mediated H3K27me3 acts to repress distinct gene sets in different cell types to maintain their specific identities. For instance, in stem cells H3K27me3 functions to maintain the stemness by repressing genes that induce cell differentiation. However, after stem cells differentiate into specific cell types, H3K27me3 instead regulates genes related to cell differentiation and maturation. Disruption of such function often leads to failed cell differentiation and tumorigenesis (Min et al., 2010). In the nervous system, knocking out *Ezh2* in neural progenitors suppresses proper progenitor cell proliferation and promotes early neuronal differentiation (Pereira et al., 2010). Lacking of EZH2 in neural progenitors also resulted in neuronal migration defects (Zhao et al., 2015). However, whether specific deletion of EZH2 in post-mitotic neurons has any effects on neural development is currently unknown, which might have distinct phenotypes from deleting EZH2 in neural stem cells. For instance, conditionally knocking out LKB1 in neural progenitors using *Emx1-Cre* led to defects in axon formation (Barnes et al., 2007), whereas knocking out LKB1 in post-mitotic neurons using *Neurod6-Cre* had no effect on axon formation but resulted in axon branching defects (Courchet et al., 2013). In this study, we used *Neurod6-Cre* mouse line to generate conditional *Ezh2* knockout mice specifically in post-mitotic neurons. Our results showed that neuronal EZH2 functioned to regulate multiple neuronal morphogenesis processes *in vivo* during development and resulted in impaired cognitive function in adult mice.

In mature neurons, a recent study showed that EZH1/EZH2 and H3K27me3 function to protect neurons from degeneration (von Schimmelmann et al., 2016). Our recent study showed that EZH2 and H3K27me3 acted to support axon regeneration (unpublished). Two previous studies have observed impaired neuronal migration when EZH2 was knocked out (Di Meglio et al., 2013) or knocked down (Zhao et al., 2015) in neural progenitors. To date, however, no study has investigated the specific *in vivo* roles of neuronal EZH2 in neural development. Our results demonstrated that in *Ezh2*^*Δ/Δ*^ mice there was impaired migration of upper layer cortical neurons at P0. In support, *in utero* electroporation of *Neurod1-Cre* plasmid into *Ezh2*^*f/f*^ mouse embryos also led to drastically decreased cortical neuron migration when examined at P0. Such neuronal migration defect was only temporary because when examined at P7, cortical neurons lacking neuronal EZH2 caught up with wild type neurons to reach the upper cortical layers. One potential mechanism is that neuronal EZH2 in early post-mitotic neurons regulates genes important for fast neuronal migration, whereas, in more mature neurons, EZH2 level decreases and plays less important roles in neuronal migration.

Our study demonstrated that deleting neuronal EZH2 resulted in more complex dendritic arborization and higher dendritic spine density *in vivo*, indicating that it acts to maintain proper dendritic arborization and dendritic spine formation during development. In contrast, our recent study showed that EZH2 was necessary for axon growth during development and regeneration (unpublished). These results suggest that during neuronal morphogenesis EZH2 functions to support axon growth and in the meantime suppress excessive dendritic development. It is likely that EZH2 regulates neuronal morphogenesis via H3K27me3-mediated gene repression. Many genes have been identified to regulate dendritic development, such as growth factors, small Rho GTPases, and cytoskeletal proteins (Brion et al., 1988; Kosik and Finch, 1987; Van Aelst and Cline, 2004). How expression of these genes is coordinated is unclear. Our study provides a potential mechanism that EZH2 coordinately regulates genes related to dendritic development. Indeed, RNA-seq comparing neurons from the control and *Ezh2*^*Δ/Δ*^ mice has revealed many genes regulated by neuronal EZH2, including *Pak3, Igf, and Bdnf*, all of which have been shown to regulate dendritic development (Dai et al., 2014; Dijkhuizen and Ghosh, 2005; Gorski et al., 2003; Pascual-Lucas et al., 2014; Schmeisser et al., 2012). We provided evidence that *Pak3* expression was repressed by EZH2-mediated H3K27me3 and blocking PAK3 function reversed increased dendritic spine density induced by EZH2 deletion. Future in depth studies of genes regulated by neuronal EZH2 will help to identify novel genes and pathways regulating *in vivo* dendritic development, providing new insights into the underlying genetic mechanisms. In this study, we also used RNA-seq to compare gene expression in post-mitotic neurons from wild type and *Nestin-Cre;Ezh2*^*f/f*^ mice, in which *Ezh2* was knocked out in neural progenitors. The identified genes partially overlap with those obtained using neurons from the *Ezh2*^*Δ/Δ*^ mice, indicating that EZH2 at different developmental stage might regulate different sets of gene expression. In support, a recent study (Durak et al., 2016) has shown that *Ezh2* expression is positively regulated by *Chd8*, a top ASD linked gene, in neural progenitors. In that study, knocking down *Chd8* resulted in down regulation of EZH2. However, upper layer cortical neurons with reduced CHD8 and EZH2 proteins show decreased complexity of dendritic arborization, which is opposite to our results. Of course, we cannot rule out the possibility that CHD8 plays an important role in the regulation of dendritic development independent of EZH2.

Dendritic arborization and dendritic spine formation are essential processes in the formation of functional circuits regulating cognitive function. In humans, mutations in *Ezh2* gene underlie the Weaver Syndrome, a genetic disease associated with intellectual disability. Both *Ezh2* and *Jmjd3* have been identified as ASD associated genes (Iossifov et al., 2012; Li et al., 2016), and *Ezh2* is a downstream target gene of *Chd8* (Durak et al., 2016). Our study revealed that specific deleting *Ezh2* in early post-mitotic neurons resulted in impaired learning and memory defects in adult mice. It is likely that changes in neuronal morphogenesis processes, especially the dendritic development, induced by neuronal *Ezh2* deletion might underlie some of the cognitive behavior phenotypes observed in patients with Weaver Syndrome or ASD. Additional more targeted cognitive behavior tests are needed in the future to further assess the roles of neuronal EZH2 in cognitive function. Previous studies have shown the roles of other epigenetic regulators in regulating dendritic development and cognitive function, such as histone deacetylase 2 (HDAC2) (Guan et al., 2009). Conversely, behavior training for learning and memory in adult mice also changes the histone modifications. For example, contextual fear conditioning increased levels of acetylation at H3 lysine 14 (H3K14), phosphorylation at H3 serine 10 (H3S10), and trimethylation at H3 lysine 4 (H3K4me3) in the hippocampus (Levenson et al. 2004; Chwang et al. 2006). Behavior training on the Morris water maze increased the acetylation at H4 lysine 12 (H4K12) and pan-acetylation of H2B (Bousiges et al. 2010). Collectively, our study provides clear and strong evidence that histone modification by EZH2 in post-mitotic neurons plays important roles in the regulation of neural developmental with cognitive consequences in adult mice.

## Methods and materials

### Animals

All experiments involving animals were performed in accordance with the animal protocol approved by the Institutional Animal Care and Use Committee of the Johns Hopkins University. The Neurod6-Cre (Nex-Cre) were generously provided by William D. Snider (Goebbels et al., 2006; Morgan-Smith et al., 2014). *Ezh2*^*f/f*^ (MMRRC_015499-UNC) mice were purchased from Mutant Mouse Resource Research Centers (MMRRC). *Ezh2*^*f/f*^ mice have been previous described (Pereira et al., 2010; Su et al., 2003). The Thy1-GFP M mice were generously provided by Richard Huganir (Duan et al., 2014). The mice have been previous described. The day of vaginal plug detection was designated as E0.5 and the day of birth as P0.

### DNA constructs and antibodies

Plasimds pCAG-dsRed was bought from addgene (Plasmid #11151). *Neurod1-Cre* was provided by Dr Franck Polleux (Columbia University). *Pak3*^K297R^ was gifted from Dr. Rick Horwitz (University of Virginia). PAK3 mutant was subcloned into pCMVmyc vector.

The following primary antibodies were used in this study: anti-EZH2 1:500 (BD biosciences, Cat.no. 612666), anti-EZH2 1:1000 (Cell signaling, Cat.no. #5246), anti-GAPDH 1:5000 (Sigma, Cat.no. G8795), anti-GFP 1:1200 (Thermo Fisher scientific, Cat.no. A10262), anti-TUJ1 1:1000 (Biolegend, Cat.no. 801201) anti-H3K27me3 1:2000 (Millipore, Cat.no. 07449), anti-H3 1:1000 (Cell signaling, Cat.no. #4499), anti-dsRED 1:1000 (Clontech, Cat.no. 632496), anti-Tbr1 1:1000 (Abcam, Cat.no. ab31940), anti-Ctip2 1:500 (Abcam, Cat.no. ab18465), anti-Foxp1 1:1000 (Abcam, Cat.no. ab16645), anti-Cre 1:1000 (Millipore, Cat.no. MAB3120).

### Primary neuronal cultures and treatment

All hippocampi and cortices were dissected in ice cold Ca^2+^- and Mg^2+^-free Hank’s balanced salt solution (HBSS; Gibco) and incubated in Papain at 37 °C for 15 min. Dissociated cells were plated on poly-D-lysine (Sigma)-coated 12mm coverslips. Cells were cultured in Neurobasal medium (Gibco) containing 2% B27 supplement (Gibco) and 50 U ml-1 penicillin, 50 µg/ml streptomycin (Gibco), 2 mM GlutaMAX (Gibco), and grown for 5 days, or 18 days with media changes every other day on glass coverslips. Plasmids pCAG-dsRed and *Pak3*^K297R^ transfection was performed at 7 d in vitro (DIV) with 1 µg of DNA and 1.5 µl of Lipofectamine (Invitrogen), as described in manufacturer’s manual (Invitrogen). After transfection, the culture medium was switched to the neuronal culture medium plus 2% FBS (Invitrogen) with medium changes every other day. Neurons were fixed in 4% paraformaldehyde for 20 min, and processed for immunocytochemistry.

### Golgi Staining

Freshly dissected mouse brains were incubated in Golgi solution A and B (FD Rapid GolgiStain Kit, FD NeuroTechnologies) for 10 days. After incubation all brains were washed thoroughly with Solution C for 72 h at room temperature, and then mouse brains were blocked and embedded in OCT embedding medium (Tissue-Tek). Coronal sections (100 mm) through the somatosensory cortex and medial CA1 were cut with a Microm HM 550 cryostat and mounted on 3% gelatin-coated slides. Staining procedures were followed as described (FD NeuroTechnologies), and slides were dehydrated in ethanol and mounted with Permount (Fisher Scientific) for microscopy. Layer II–III neurons from the somatosensory cortex were included in our analyses.

### Immunohistochemistry and fluorescence intensity quantification

Immunohistochemistry of cultured neurons and brains were performed and quantified using standard methods. In brief, the cultured cells were washed with PBS (Invitrogen), and then fixed in 4% PFA. Neurons were blocked in 1% BSA (in 0.2% PBST), incubated with primary antibodies overnight at 4°C, and visualized with secondary antibodies.

Perfused brains were processed and sections were stained as described (Cubelos et al., 2008). Brains were gained and post-fixed in 4% PFA for 24 h at 4°C, and then followed by 15%, 30% (w/v) sucrose in PBS at 4°C until brains sunk. Brains were sectioned in from 40 to 50μm. Sections were incubated with primary antibodies overnight at 4°C, and incubated with secondary antibodies for 1h at room temperature.

All the images were taken by Zeiss 510 confocal microscopy. Fluorescence intensity were quantified by Image J software ((NIH, available from http://rsb.info.nih.gov/ij). First, the fluorescence intensity from each image stacks was subtracted by their background. Next, the average fluorescence per cell was calculated by measuring their average mean values.

### Western blot analysis and quantification

Tissue lysate were run on NuPAGE Novex 4-12% Bis-Tris Protein Gels (Thermo Fisher Scientific). Western blotting were performed as described (Saijilafu et al., 2013). Novex gels were run at 150 constant voltage to separate them, and transferred onto Immu-blot membranes (Bio-Rad) at a constant voltage (35V for 3hours). Membranes were blocked using 5% milk prepared in TBS-T (50 mm Tris-HCl (pH 7.4), 150 mm NaCl, 0.1% Tween-20) for 1 h at room temperature. Membranes were incubated with the primary antibodies overnight at 4 °C. Secondary antibodies conjugated with horseradish peroxidase (Thermo Fisher Scientific) were incubated for 1 h at room temperature. Following washing with TBS-T, immunoreactivity signals were detected by enhanced chemiluminescence (Perkin-Elmer). Western blot were quantified by Image J with standard method.

### Morphological Analysis

All images were captured with a Zeiss 510 laser scanning confocal microscope. Dendritic processes, spine number, and spine morphology of individual neurons of the somatosensory cortex and hippocampus were measured with ImagJ or Imaris.

Dendrites arising from the cell body were considered as first-order segments until they bifurcated symmetrically into second-order segments; dendritic branches arising from the first-order segments were considered as second-order segments until they bifurcated symmetrically into third-order segments. The following parameters for each reconstructed cortical neuron were analyzed: (i) total dendritic branch length, representing the summed length of dendritic segments; (ii) total dendritic branch tip number (TDBTN), representing the total number of dendritic segments; (iii) average dendritic branch length, representing the individual dendritic branch length.

Spine density was calculated by quantifying the number of spines per measured length of the parent dendrite and expressed as the number of spines per 10-µm length of dendrite. The length of each dendritic segment used for spine densitometry was around 20 µm. For each neuron, spines in the second-order branches were quantified in P28-P30 mice All spine density quantification in *in vitro* culture was obtained from a 20 µm dendrite segment at the second-order branches of neurons.

### Dendrite branching analysis and Sholl analysis

All GFP-positive image stacks from transfected cortical neurons were taken as described. Dendrite branching was calculated manually by tracing dendrites of neurons. Sholl analysis was performed by drawing concentric circles centered on the cell soma using ImageJ. The starting radius was 5 µm and the interval between consecutive radii was 5 µm.

### Behavioral studies

All behavioral experiments were performed on 3-4 month old *Ezh2*^*Δ/Δ*^ mice and their *Ezh2*^*flox/flox*^(control) littermates. Both male and female mice were included.

#### Y-Maze

The y-maze test was used to assess working memory and short-term spatial recognition memory in mice. To test for working memory, spontaneous alternations were assessed. Mice were placed at the end of one of the three arms of the maze (San Diego Instruments, San Diego, CA]) and allowed to explore the maze for 5 minutes. Videos were recorded and analyzed by Top Scan software (CleverSys Inc., Reston, VA). Spontaneous alternations were measured as percentage of correct sequences of successive, non-repeated visits to all three arms. 5 days later mice were tested again on the y-maze for recognition memory. For trial 1, one of the three arms was blocked and the animals were allowed to explore the other two arms for 5 minutes. 12-15 minutes later, animals were returned to the maze with no arms blocked and allowed to explore for 5 minutes. Time spent in each arm was quantified, with more than 33% of time spent in the previously blocked arm was identified as recognition memory. Time spent in the blocked arm was compared between the experimental and control mice.

#### Elevated Plus Maze

To test anxiety of the mice, elevated plus maze (EPM) was performed. Mice were placed in the center of a t shaped maze (San Diego Instruments, San Diego, CA) that had two open arms and two arms enclosed by the walls. Mice were allowed to EPM sfor 5 minutes. Videos of the mice movements were recorded and analyzed for % of time spent in the open and closed arms by Top Scan software (CleverSys Inc., Reston, VA).

#### Novel Object Recognition

Object recognition memory was tested in the mice using novel object recognition test. For 3 days prior to testing, mice were allowed to habituate to custom made 25 cm x 25cm c 25cm acrylic boxes for 10 minutes. On the 4^th^ day, two identical objects were placed in the center of the boxed equidistance from each other and the walls and mice were allowed to explore the objects for 10 minutes. One hour later, one of the two identical objects was replaced with a novel object. Mice were returned to the box and were allowed to explore the familiar and novel object. Time spent sniffing the familiar and novel object was automatically measured with the Top Scan software (CleverSys Inc., Reston, VA).

#### Open field test

General locomotor activity was evaluated in the activity chambers (SDI, San Diego, USA) for 30 minutes. The time spent in the center and periphery of the arena as well as number of rears were automatically assessed.

### In utero electroporation

All pregnant females were deeply anaesthetized with avertin via intraperitoneal injection. Plasmids were mixed with fast green and microinjected into the lateral ventricle of embryos using a picospritzer. Embryos were exposed at E15.5. Embryos were electroporated with five pulses of 35 V (50 ms on, 950 ms off at 1 Hz) through 5-mm tweezer electrodes connected to a square wave electroporator (CUY21, π Protech).

### Chromatin immunoprecipitation (ChIP) assay

ChIP assays were performed with a commercial kit (Chip-IT kit, Active motif). The cortices from P2 wildtype mice were minced and crosslinked in 1% formaldehyde (F8775, Sigma) for 15 min and the reaction was stopped by adding glycine (0.125 M). Nuclei of the cells were precipitated, lysated, and sonicated on ice 20 times for 5 s (duty cycle 40%, microtip limit 4) (Vibra-Cell V 50, Sonics Materials) (average fragment size of 400 bp). Part of supernatant was saved as input. The antibody used was a polyclonal anti-H3K27me3 (Millipore) and an unrelated rabbit IgG. Immunoprecipitates were mixed with protein G magnetic beads and incubated overnight at 4°C and washed, and protein/DNA complexes were eluted with cross-links reversed by incubating in ChIP elution buffer plus proteinase K for 2 h at 65°C. DNA was purified using spin columns and analyzed in duplicate by Q-PCR. Primer sequences for Q-PCR are described in Supplemental tables. Fold enrichment is expressed as the ratio of H3K27me3 signal to IgG signal.

### Real-time PCR

Total RNA from control and knockout mice was extracted using Trizol reagent (Invitrogen). RNA samples were treated with DNase I (Invitrogen) and reverse transcription was performed using transcriptor first strand cDNA synthesis kit (Roche). The LightCycler 480 SYBR Green I master mix (Roche) was used for real-time PCR. Threshold cycle (Ct) was determined on the linear phase. Relative gene expression fold difference was calculated by 2^−Δnormalized Ct^. PCR primers are listed in Supplementary Table.

### Electrophysiology

Whole-cell electrophysiology and data analysis. Voltage-clamp wholecell recordings were obtained from cultured neurons at room temperature (22–25°C). An external solution containing 150 mM NaCl, 3 mM KCl, 10 mM HEPES, 6 mM mannitol, 1.5 mM MgCl2, and 2.5 mM CaCl2, pH 7.4, was used for the recordings. Glass pipettes with a resistance of 5– 8 mΩ were filled with an internal solution consisting of 110 mM cesium gluconate, 30 mM CsCl, 5 mM HEPES, 4 mM NaCl, 0.5 CaCl2,2mM MgCl2,5mM BAPTA, 2 mM Na2ATP, and 0.3 mM Na2GTP, pH 7.35. Neurons were held at a holding potential of −70 mV for mEPSCs. For recording mEPSCs, 1 M tetrodotoxin and 100 M picrotoxin were added to the external recording solution. The signal was filtered at 2 kHz and digitized at 20 kHz using an Axon Digidata 1440A analog-to-digital board (Molecular Devices). Recordings were performed from three independent cultures. Recordings with a pipette access resistance of 20 m and with 20% changes during the duration of recording were included. The mEPSC recordings was analyzed using Mini Analysis software (Synaposoft) with an amplitude threshold set at 6 – 8 pA. The frequency, amplitude, and decay were measured in each group.

### Genome-wide RNA-sequencing and bioinformatic analyses

Cortical neurons from either *Nestin*-Cre; *Ezh2*^*f/f*^ *or Neurod6*-Cre; *Ezh2*^*f/*f^ and their control litter mates were isolated and cultured for 3 days. Following total RNA was isolated using RNeasy Plus Kit (Qiagen), RNA was quality controlled and quantified using an Agilent 2100 Bioanalyzer. High-throughput sequencing was performed using the Illumina HiSeq 2000 platform at the JHMI deep sequencing & microarray core.

The data analysis of RNA-seq used the following workflows. First, transcript isoform level analysis was performed by aligning the 54 base cDNA reads against the reference genome mm9 build using a Burrows-Wheeler transform based short read aligner (BWA), and the aligned reads were visualized using Integrated Genome Viewer. The read alignment files were imported into Partek Genomics Suite (Partek® Genomics SuiteTM) and RPKM (reads per kilobase of exon model per million mapped reads) counts for each of the 28,157 transcripts defined in the UCSC refflat annotation file were calculated. A stringent filtering criterion with RPKM value of 1.0 was used to obtain transcripts. The RPKM values of filtered transcripts were log-transformed using log2 (RPKM + offset) with an offset value of 1.0. Fold changes in transcript expression and *p*-values were computed using ANOVA, and significantly altered transcripts were selected by applying fold change-cutoff of 2. Alternatively, gene level analysis was performed by aligning the filtered reads to the mouse reference genome build mm9 using the ELAND (Efficient Local Alignment of Nucleotide Data) algorithm (Anthony J Cox, Solexa Ltd Saffron, Walden, UK). First, raw and RPKM normalized counts were calculated for gene models (as defined in <u>UCSC RefGene</u>). Subsequently, we applied SAM (significance analysis of microarrays) to gene level RPKM to identify RNAs with a fold change greater than 2 and false discovery rate (FDR) of less than 25%. Briefly, the boundaries of exons were obtained from the RefGene database (<u>Genome</u>), and the numbers of mapped reads on each exon were calculated. To compare to human data sets, mouse gene names were converted to human homologs using MGI annotation database (http://www.informatics.jax.org/homology.shtml). Gene ontology (biological process) was assessed using Metascape (http://metascape.org/gp/index.html#/main/step1) web servers on substantially expressed genes, meaning minimal log2(RPKM) of 1 among upregulated genes upon EZH2 deletion, or minimal log2(RPKM) of 1 among downregulated genes in the control.

### Statistics

All plots were generated and statistical analyses were performed using Graphpad Prism 5.0 software. Results are presented as mean ± SEM. Sample size was not predetermined but numbers of samples are consistent with previous publications. Two-tailed t-tests were used for comparison of two data sets. Equal variances between groups and normal distributions of data were presumed but not formally tested. Molecular and biochemical analyses were performed using a minimum of three biological replicates per condition. Behavioral experiments require larger data sets due to increased variability.

## Acknowledgements

We thank Randal Hand for the Neurod1-Cre plasmid, and Rick Horwitz for the PAK3^K297R^ plasmid. We thank Zhongxian Jiao for mouse husbandry and Michele Pucak for technical assistance in confocal microscopy. The study was supported by grants (to F.Q.Z.) from NIH (R01NS064288, R01NS085176, R01GM111514, R01EY027347), the Craig H. Neilsen Foundation (259450), and the BrightFocus Foundation (G2017037).

## Author Contributions

M.Z., C-M. L. and F-Q. Z. conceived the study and designed the project; M.Z. performed most of the experiments; Y.Z. and R.L.H. contributed to the in utero-electroporation experiment; Q.X., X.D., E.L. and G.D. contributed to the electrophysiological experiments; M.Z., C-M. L, J.C., and M.V.P. contributed to the mouse behavior experiments and data analysis; M.Z. and P.H. analyzed the data. G-H.W. and J.Q. analyzed the RNA-Seq data; M.Z. and F-Q.Z. wrote the manuscript with contribution from all authors.

## Supplementary figure legends

**Supplementary Figure S1.**
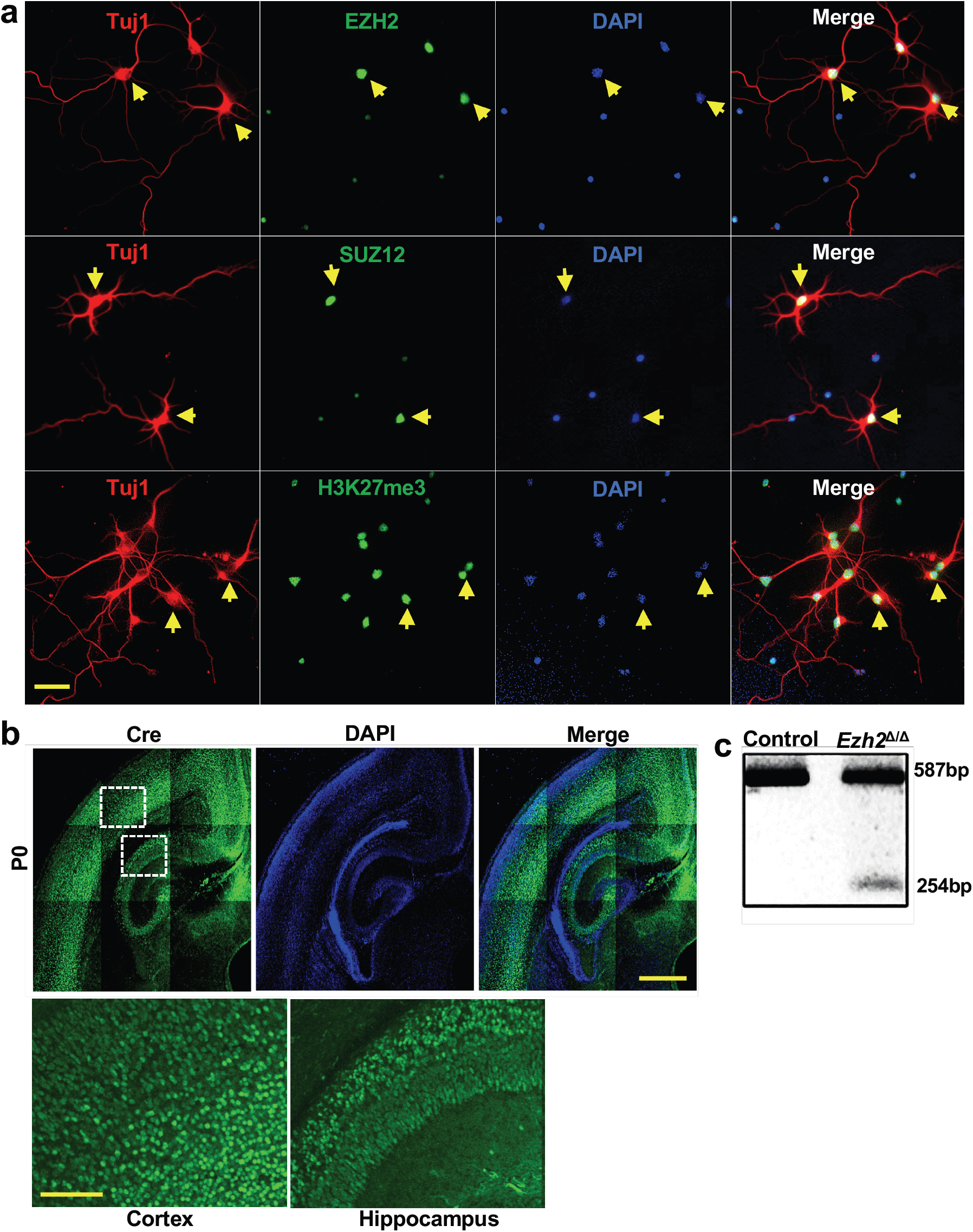
Expression of EZH2, SUZ12, and H3K27me3 in cultured excitatory neurons. (a) Representative images showing the expression of EZH2, SUZ12 and H3K27me3 in cultured primary cortical neurons. Yellow arrowheads indicate Tuj1 positive neurons. Scale bar, 50µm. (b) Top: representative images of P0 mouse coronal brain section stained with anti-Cre antibody showing the expression of Cre at cortical and hippocampal region. Bottom: the enlarged images of two white dashed boxes shown in the top panel. Scale bar, 500µm in the bottom panel and 2 mm in the top panel. (c) NeuroD6-Cre mediated recombination in the adult mouse brain was confirmed by RT-PCR. Representative PCR band image showing a 254bp fragment generated by NeuroD6-Cre mediated *Ezh2* deletion.

**Supplementary Figure S2.**
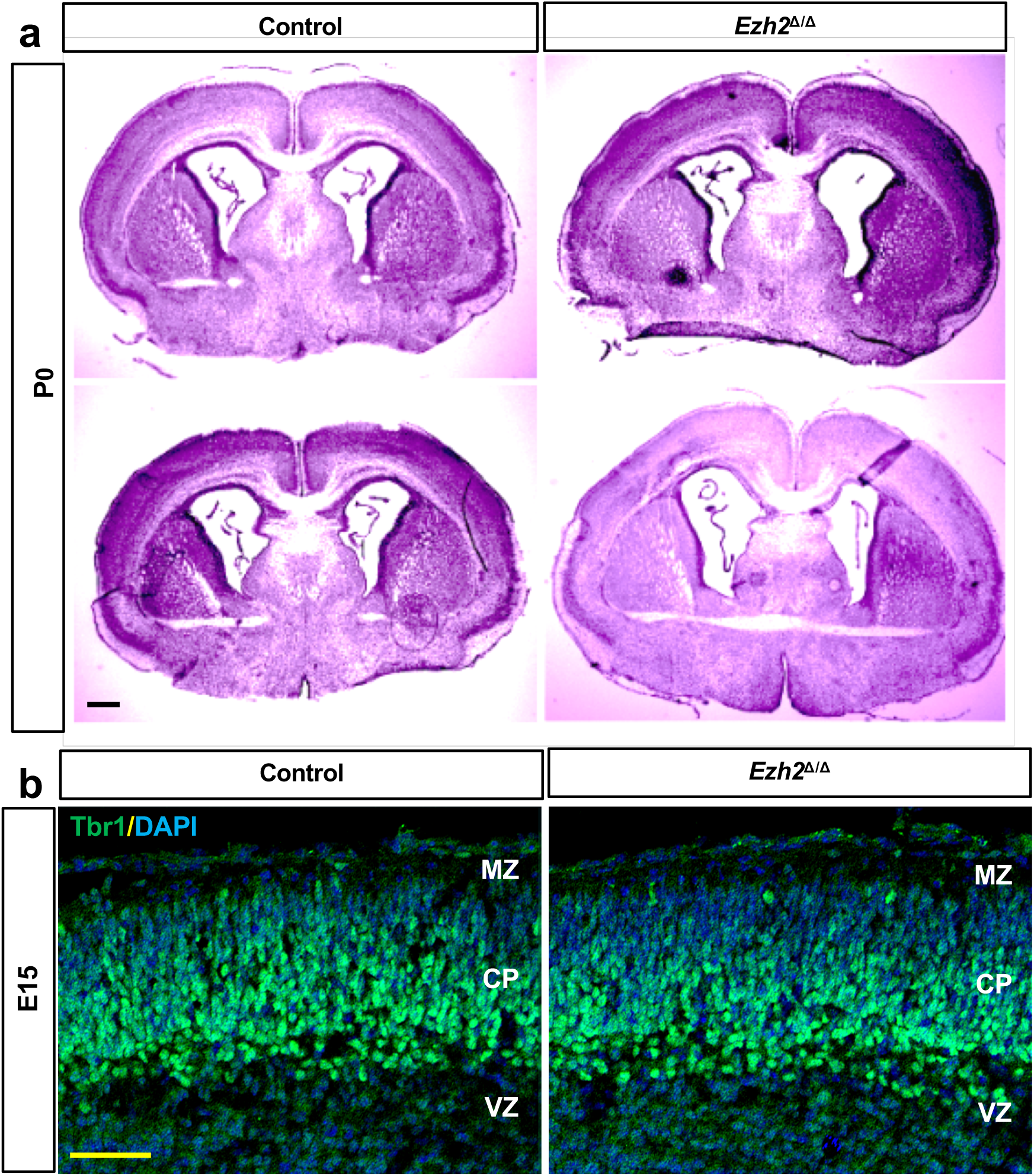
Deletion of EZH2 manifested no structural brain abnormalities. (a) Representative nissl staining images of coronal sections from P0 control and *Ezh2*^Δ/Δ^ mice showing no obvious difference between the control and the *Ezh2*^Δ/Δ^ mice. (b) Representative images of coronal brain sections stained with anti-Tbr1 antibody and DAPI showing that *Ezh2*^Δ/Δ^ mice have normal early neurogenesis at E14.5. Scale bar, 100µm

**Supplementary Figure S3.**
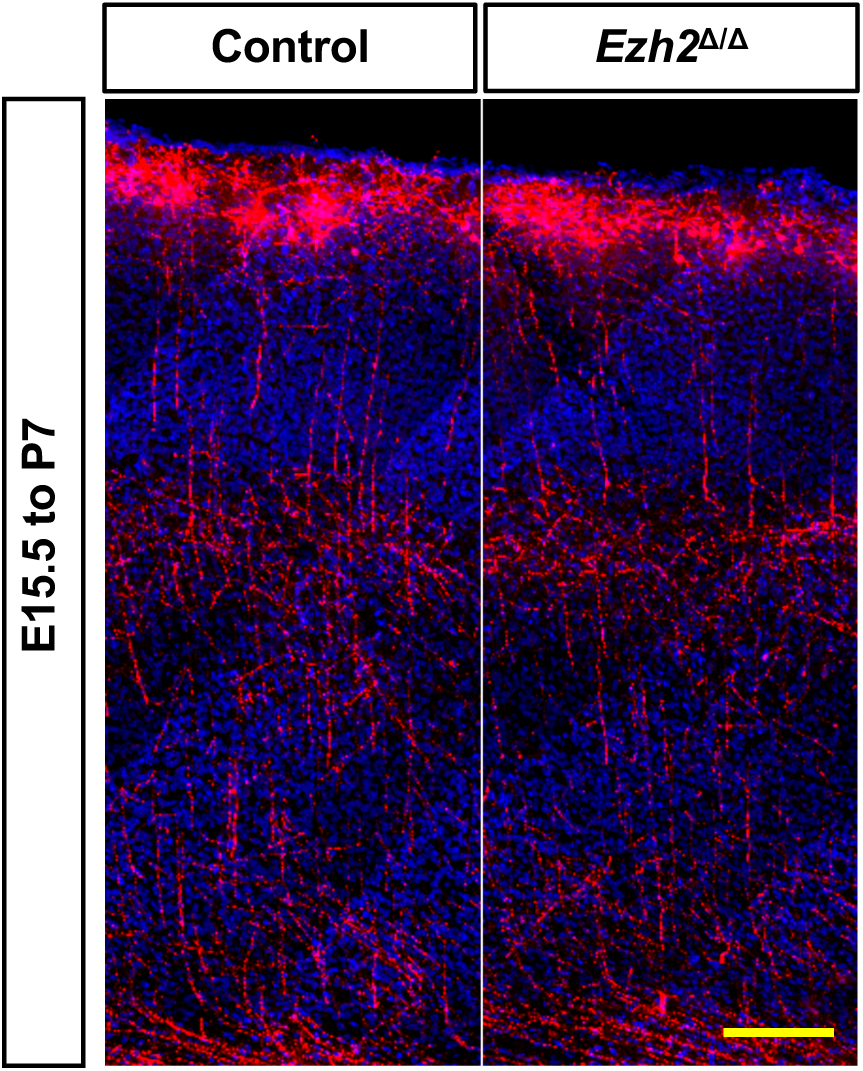
Lack of EZH2 did not impair the cortical neuronal migration on P7. Representative confocal images of mouse cortices in utero electroporated with dsRED or dsRED/NeuroD-cre. The electroporation was performed at E15 and the pups were harvest at P7.

**Supplementary Figure S4.**
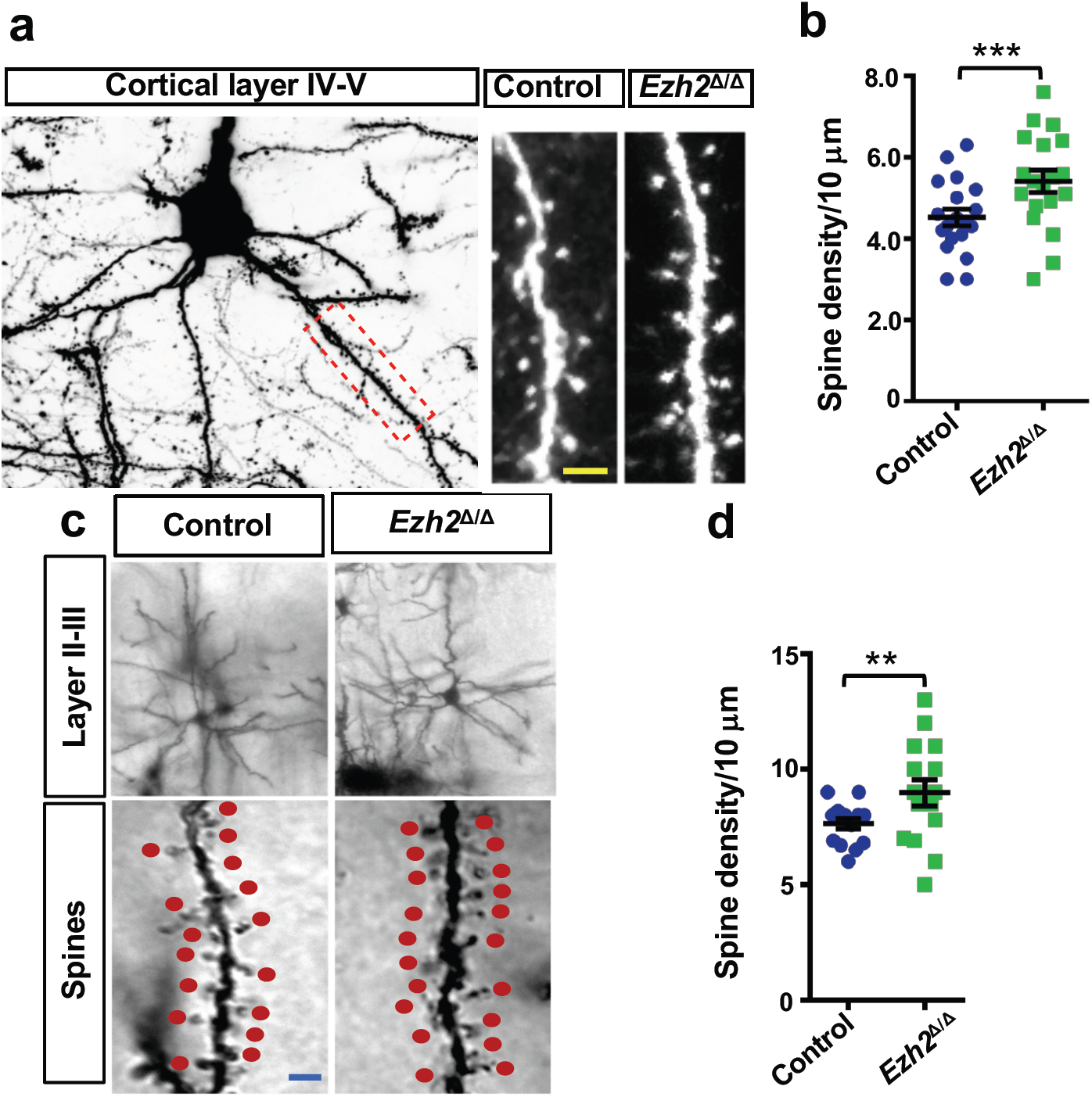
Spine density was increased in somatosensory cortex of *EZH2*^Δ/Δ^ animals. (a) Representative images of increased spine density in basal region of layer IV-V neurons from *Ezh2*^*f/f*^; Thy1-GFP and *Ezh2*^*Δ/Δ*^; Thy1-GFP mice. (b) Quantification analysis of spine density in deep region of cortical neurons from control and *Ezh2*^*Δ/Δ*^ mice. n=25 for control mice and n=26 for *Ezh2*^*Δ/Δ*^ mice, *P*=0.0002. (c) Representative images of increased spine density in basal region of layer II-III neurons from *Ezh2*^*f/f*^; and *Ezh2*^*Δ/Δ*^ mice labeled with Golgi staining. (d) Quantification analysis of spine density in layer II-III region of cortical neurons from control and *Ezh2*^*Δ/Δ*^ mice. n=25 for control mice and n=26 for *Ezh2*^*Δ/Δ*^ mice, *P*=0.0095. Data are presented as mean ± SEM. n.s., no significant difference, \**P*<0.05, \*\**P*<0.01, \*\*\**P*<0.001, \*\*\*\**P* < 0.0001, compared to control if not designated.

**Supplementary Figure S5.**
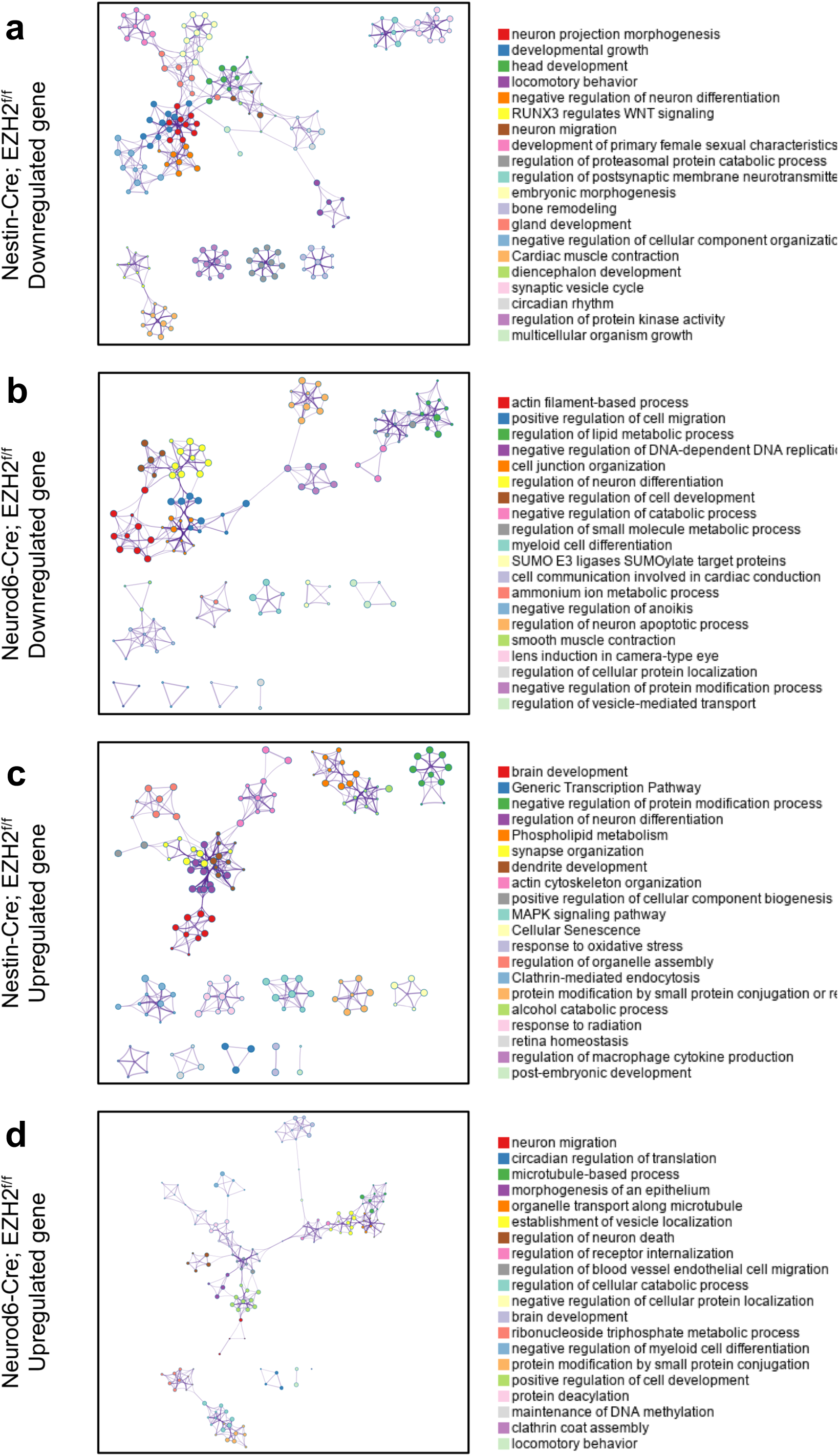
Network of enriched gene ontology (GO) terms among each differential gene set depicts connections among GO terms. Subgroups of enriched GO terms shown in Figure 6 have been plotted and visualized using the Metascape (http://metascape.org) and Cytoscape5. The program constructs GO term network among terms with a similarity > 0.3, which are connected by edges. Each node represents one enriched GO term, where node size is the number of genes within a GO term. GO terms within the same cluster are visualized with the same color. *P*-values of the enrichments refer back to the Figure 6, which shows maximal level of p-value is 1.0E-3. (a) GO term network of downregulated genes in the Nestin-Cre;EZH2^f/f^ compared with the control. (b) GO term network of downregulated genes in the Neurod6 (Nex)-Cre;EZH2^f/f^ compared with the control. (c) GO term network of upregulated genes in the Nestin-Cre;EZH2^f/f^ compared with the control. (d) GO term network of upregulated genes in the Neurod6 (Nex)-Cre;EZH2^f/f^ compared with the control.

**Supplementary Figure S6.**
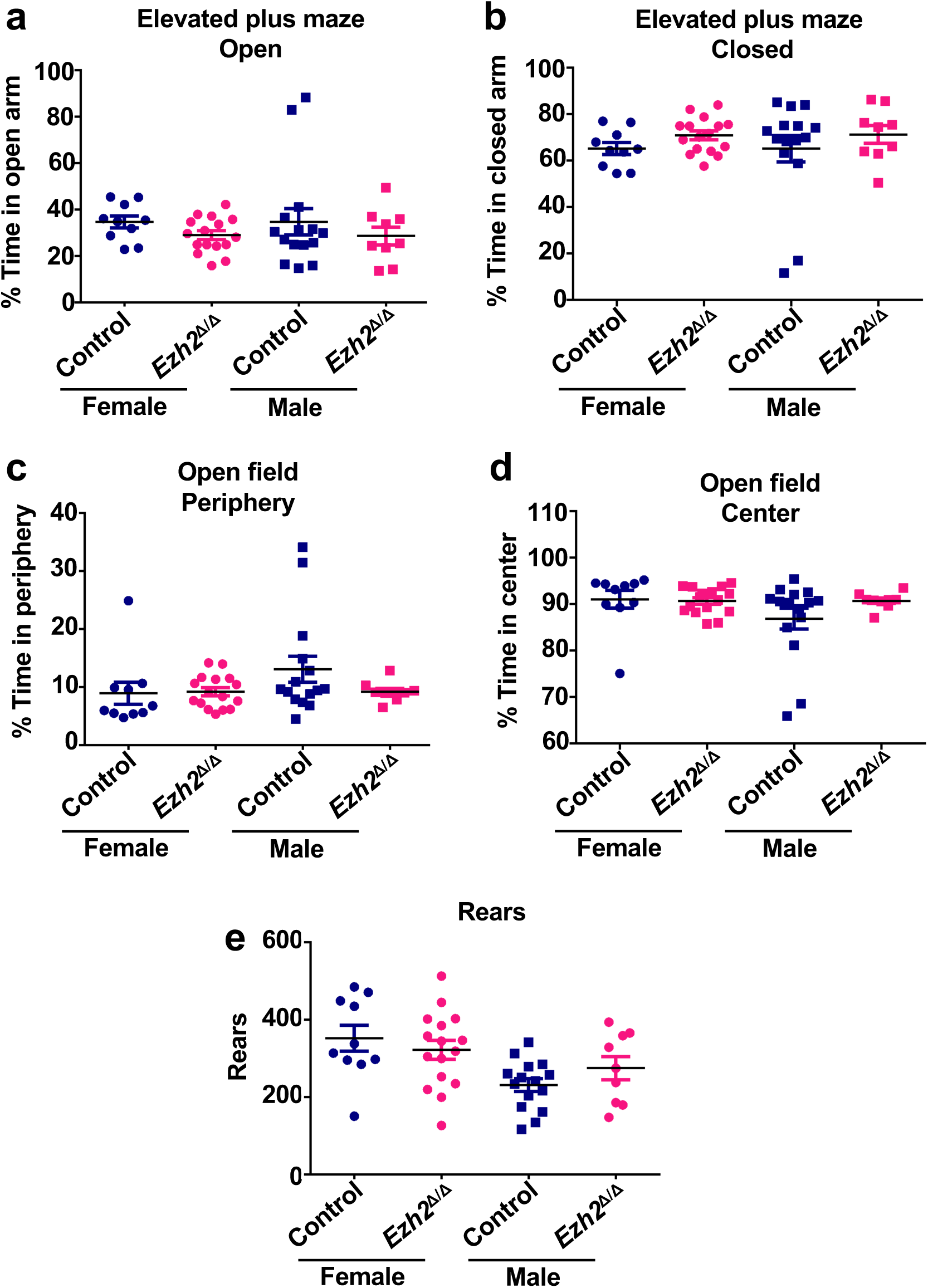
*EZH2*^*Δ/Δ*^ mutant mice show little defects in elevated plus maze test and the open field test. (a) Quantification of time spent in the open arms of the elevated plus maze. Female: n=10 mice for control mice and n=16 for *Ezh2*^*Δ/Δ*^ mice, *P*=0.6808; male: n=15 for control mice and n=9 for *Ezh2*^*Δ/Δ*^ mice, *P*=0.6093. (b) Quantification of time spent in the closed arms of the elevated plus maze. Female: n=10 for the control group and n=16 for the *Ezh2*^*Δ/Δ*^ group, *P* =0.6808; male: n=15 for control mice and n=9 for *Ezh2*^*Δ/Δ*^ mice, *P* =0.6093. (c-e) Quantification of alterations in time spent peripheral zone (c), central zone (d), and the rearing behavior (e). Female: n=10 for control mice and n=16 for *Ezh2*^*Δ/Δ*^ mice; male: n=15 for control mice and n=9 for *Ezh2*^*Δ/Δ*^ mice. Open field peripheral: *P*=0.8551 for female mice and *P* =0.2052 for male mice. Open field center: *P*=0.8561 for female mice and *P*=0.3643 for male mice. Open field rearing: *P* =0.4703 for female mice and *P* = 0.1854 for male mice. Data are presented as mean ± SEM. n.s., no significant difference, \**P*<0.05, \*\**P*<0.01, \*\*\**P*<0.001, \*\*\*\**P* < 0.0001, compared to control if not designated.

